# Standardized quality control workflow to evaluate the reproducibility and differentiation potential of human iPSCs into neurons

**DOI:** 10.1101/2021.01.13.426620

**Authors:** Carol X.-Q. Chen, Narges Abdian, Gilles Maussion, Rhalena A. Thomas, Iveta Demirova, Eddie Cai, Mahdieh Tabatabaei, Lenore K. Beitel, Jason Karamchandani, Edward A. Fon, Thomas M. Durcan

## Abstract

Induced pluripotent stem cells (iPSCs) derived from human somatic cells have created new opportunities to generate disease-relevant cells. Thus, as the use of patient-derived stem cells has become more widespread, having a workflow to monitor each line is critical. This ensures iPSCs pass a suite of quality control measures, promoting reproducibility across experiments and between labs. With this in mind, we established a multistep workflow to assess our newly generated iPSCs for variations and reproducibility relative to each other and iPSCs obtained from external sources. Our benchmarks for evaluating iPSCs include examining iPSC morphology and proliferation in two different media conditions and evaluating their ability to differentiate into each of the three germ layers, with a particular focus on neurons. Genomic integrity in the human iPSCs was analyzed by G-band karyotyping and a qPCR-based test for the detection of hotspot mutations test. Cell-line identity was authenticated by Short Tandem Repeat (STR) analysis. Using standardized dual SMAD inhibition methods, all iPSC lines gave rise to neural progenitors that could subsequently be differentiated into cortical neurons. Neural differentiation was analyzed qualitatively by immunocytochemistry and quantitatively by qPCR for progenitor, neuronal, cortical, and glial markers. Taken together, we present a multistep quality control workflow to evaluate variability and reproducibility across and between iPSCs.

## 1. Introduction

Human pluripotent stem cells can give rise to any cell type when exposed to the appropriate developmental cues, holding enormous potential for tissue engineering, regenerative medicine and disease modeling. The growing number of iPSC lines and NIH-registered human embryonic stem cell (ESC) lines ensures that patient-derived pluripotent stem cells are now readily available to researchers, helping to accelerate our understanding of biology and disease and the development of therapies across disease areas [1]. These developments underscore the need for iPSC quality standards that are sufficiently stringent to ensure that findings can be compared and results reproduced across laboratories [2, 3].

For instance, it is critical that the growth parameters of a given iPSC cell-line are optimized. Since the advent of iPSC technology, culture conditions have become more standardized, helping to increase reproducibility between and within cell-lines. Several media formulations have been developed to grow and maintain iPSCs [4, 5]. Thus, depending on the media used, iPSC growth can vary, making it imperative that the optimal growth media and maintenance conditions be defined before working with any new cell-line and to reduce variation between lines [5–8]. Moreover, numerous studies on pluripotent ESCs and iPSCs have identified genomic integrity as a critical issue when monitoring the quality of cells for use in research or clinical applications. Common alternations that arise in the genome during iPSC reprogramming, from culture conditions [9] or off-target CRISPR/Cas9 genome editing [10, 11] can include, but are not limited to, (1) chromosomal/karyotypic abnormalities [12–14] (2) copy number variations (CNVs), (3) small genomic insertions and deletions, or (4) single nucleotide variants [15]. When present, these genomic abnormalities often alter the biological properties of hiPSC-derived models. Thus, before a given iPSC is used, it should be tested for alterations at the genomic level [3, 16, 17].

Nonetheless, each benchmark for evaluating iPSCs has its own strengths and limitations in terms of sensitivity, costs, time and effort. Considering these points, we have developed a rigorous quality control workflow to evaluate newly generated iPSC lines. In this paper, we present a workflow combining these quality control (QC) assays that we used to test for variability across ten newly generated hiPSC lines derived from healthy individuals, relative to existing commercial lines. With these cell-lines, we established a workflow to monitor hiPSC morphology and proliferation in two different media. As well, the iPSCs were authenticated through Short Tandem Repeat (STR) profiling, to verify that each iPSC line generated matched the parental cell from which it was reprogrammed. Genomic integrity in the hiPSCs was analyzed by G-band karyotyping and qPCR-based profiling for genomic hotspot regions that are commonly altered during reprogramming [18–20]. We also evaluated the pluripotency of our lines by examining their ability to form embryoid bodies (EBs), to differentiate into each of the three germ layers, and their potential to form cortical neurons. We focused on cortical neurons, as methods to generate these neurons from iPSCs are well established, and represent a cell type broadly studied across disease areas of the brain, from Alzheimer’s disease to neurodevelopmental disorders [21, 22]. While all iPSC lines were capable of generating cortical neurons under standard differentiation protocols [23], line-to-line variability in the number of neurons formed, and their overall morphology was observed, highlighting the importance of deep phenotyping of cell lines before they are used in a given downstream application. Taken together, we developed a multistep, QC workflow (**Figure 1**) to validate our newly generated iPSCs lines and their ability to reproducibly differentiate into human cortical neurons for downstream applications. Ensuring iPSC colonies maintain an undifferentiated morphology, determining the growth rate of each line, and assessing the pluripotency and differentiation potential of each line are essential quality control measures, to reduce variability across different iPSCs.

**Figure 1.**
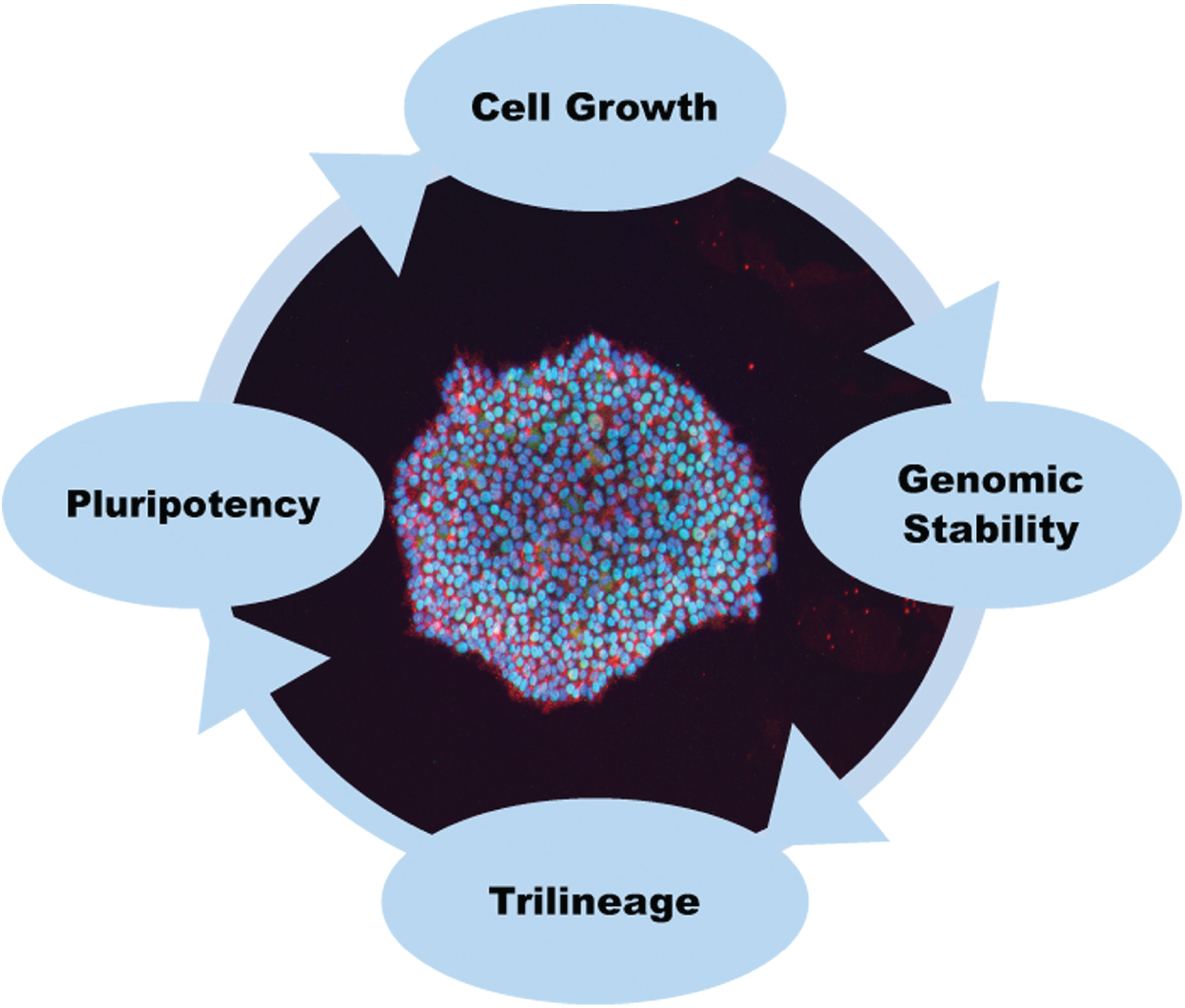
Multistep workflow for phenotyping of hiPSCs. Schematic representation of a multistep QC workflow to monitor the morphology and proliferation of newly generated iPSCs, genomic integrity, pluripotency, and an ability to form cells of the three germ layers.

## 2. Materials and methods

### 2.1. Cell lines

The lines used were: H9 hESC (Wicell), NCRM1 (NIH), KYOU-DXR0109B (ATCC), AIW002-02, AJC001-5, AJG001-C4 (same donor but reprogrammed using different methods or from different cell types), AJD002-3, TD03 (same donor but reprogrammed with different methods), TD2, TD10, TD22, 3448 and 3450. The complete profiles of the iPSCs are listed in **Table 1**. The use of iPSCs and stem cells in this research was approved by the McGill University Health Centre Research Ethics Board (DURCAN_IPSC / 2019-5374).

**Table 1:** Overview of hiPSCs.

| Cell line | Donor | Cell Type | Donor age | Sex<sup>1</sup> | Cell source | Reprogramming | Media<sup>2</sup> |
| --- | --- | --- | --- | --- | --- | --- | --- |
| H9 |  | ESC | NA | F | WiCell |  | mTeSR1 |
| NCRM1 | A | cord blood | NA | M | NIH | Episomal | mTeSR1 |
| KYOU-DXR0109B | B | fibroblast | 36 | F | ATCC | Sendai virus | mTeSR1 |
| AIW002-02 | C | PBMC | 37 | M | The Neuro | Sendai virus | mTeSR1 |
| AJC001-5 | C | fibroblast | 37 | M | The Neuro | Sendai virus | mTeSR1 |
| AJG001-C4 | C | PBMC | 37 | M | The Neuro | Episomal | mTeSR1 |
| TD02 | D | PBMC | 48 | F | The Neuro | Episomal | mTeSR1 |
| 3448 | E | PBMC | 48 | M | The Neuro | Episomal | E8 |
| 3450 | F | PBMC | 37 | M | The Neuro | Episomal | E8 |
| AJD002-3 | G | PBMC | 44 | M | The Neuro | Sendai virus | E8 |
| TD03 | G | PBMC | 44 | M | The Neuro | Episomal | E8 |
| TD10 | H | PBMC | 64 | F | The Neuro | Episomal | E8 |
| TD22 | I | PBMC | 59 | M | The Neuro | Episomal | E8 |

<sup>1</sup> M, male; F, female; <sup>2</sup> optimal media for cell growth
The passage number of NEURO hiPSC lines are 8-12; H9: P30+P5; NCRM1: P50+P11; KYOU-DXR0109B: +P3 after cell had been purchased from ATCC, no passage information on product datasheet.

### 2.2. iPSC reprogramming

PBMCs and skin fibroblast cells were obtained through the C-BIG Repository at the Montreal Neurological Institute (The Neuro). AIW002-02, AJC001-5 and AJD002-3 were generated using the CytoTune™-iPS 2.0 Sendai Reprogramming Kit (iPSQuebec Platform, Laval University) and TD02, TD03, TD10 and TD22 through episomal reprogramming (Axol Biosciences, UK). AJG001-C4, 3448 and 3450 were generated in house by episomal reprogramming [24]. Briefly, PBMCs were cultured for 6 days, 2~3 x10^6^ cells were nucleofected with episomal plasmids (pEV-OCT4-2A-SOX2, pEV-MYC, pEV-KLF4, and pEV-BC-XL, a generous gift from Dr. XB Zhang, Loma Linda University). The transfected PBMCs were plated on mitomycin C-treated mouse embryonic fibroblasts cultured in KnockOut DMEM/F12 supplement with 20% Knockout serum supplement, 50 ng/ml fibroblast growth factor 2, 1x Insulin-Transferrin-Selenium and 50 mg/ml 2-phospho-L-ascorbic acid. The cultures were refreshed with 2 ml of the above medium every 2 day until day 8. When colonies displayed iPSC morphology (between day 6-8 post-transfection), cells were fed with mTeSR1 medium (Stemcell Technologies) supplemented with 0.25 mM sodium butyrate every 2 days until day 14. Colonies were picked manually on day 14-16, and cultured on Matrigel-coated dishes every 5–7 days until after 5 passages when they were cryopreserved for further testing and profiling.

### 2.3. Culture conditions for iPSCs

iPSCs were cultured and expanded on plates coated with Matrigel (Corning, 354277) in either mTeSR1 or E8 (ThermoFisher Scientific, A1517001) media. Cells were maintained at 37°C with 5% CO_2_ with daily media changes and split when cells reached 70-80% confluency (within 5-7 days of seeding). Any iPSC colonies with irregular borders, spontaneous differentiation or transparent centers were manually removed prior to splitting. Cells were passaged by incubation in Gentle Cell Dissociation media (Stemcell Technologies, 07174) for 4 minutes at 37°C to obtain single cells or RT for 6 minutes to obtain small aggregates of colonies. The following cell densities were used: 2×10^4^ cells/well in 24 well plates for immunocytochemistry; 2×10^4^/60 mm dish for daily morphology imaging; 2×10^5^/well in 6 well plates for EB formation.

### 2.4. Crystal violet assay

For this assay, 4000 cells/well were plated in Matrigel-coated 96-well plates with mTeSR1 or E8 media. After 2, 4 or 6 days in culture, plates were rinsed with PBS to remove non-attached cells and fixed with 3.7% PFA for 5 minutes, before staining with 0.05% crystal violet (CV, Sigma,46364) diluted in water for 30 minutes. The CV dye was thoroughly washed away using distilled water, and the plates dried at RT. Once dried, the plates were imaged and quantification performed [25] by adding 100 μl of methanol (Fisher Chemical, 32435K7) to the wells to solubilize the CV, followed by measurement of the O.D. at 540 nm (OD540) with an EnSpire Multimode Plate Reader (Perkin Elmer).

### 2.5. RNA isolation, cDNA synthesis, and qPCR analysis

RNA was purified with a NucleoSpin RNA kit (Takara) according to the manufacturer’s instructions. cDNA was generated using the iScript Reverse Transcription Supermix (BioRad). Quantitative real-time PCR was performed on the QuantStudio 5 Real-Time PCR System (Applied Biosystems) using the primers listed in **Table S1**. Raw data was processed using a custom Python script available at https://github.com/neuroeddu/Auto-qPCR. The cycle threshold (CT) values for technical triplicates were tested for outliers. Relative gene expression was calculated by using the Comparative CT Method (ΔΔCT method), where the endogenous controls were GADPH or ACTB expression. The reference sample varied by experiment and is indicated in each plot.

### 2.6. Short Tandem Repeat (STR) analysis

All newly generated iPSCs were authenticated through STR analysis with the GenePrint® 10 System (Promega, B9510) at The Centre for Applied Genomics, the Hospital for Sick Children, Toronto. Briefly, genomic DNA from the iPSCs or the source material for the iPSCs, which in this case was PBMCs, was extracted with a Genomic DNA Mini Kit (Blood/Cultured Cell) (Geneaid, GB100). Ten ng of genomic DNA was mixed with GenePrint® 10 primer pair mix to permit co-amplification and detection of ten human loci, including all ASN-0002-2011 loci (TH01, TPOX, vWA, CSF1PO, D16S539, D7S820, D13S317, D5S818) plus Amelogenin for gender identification and one mouse locus D21S11. These loci collectively provide a genetic profile with a random match probability of 1 in 2.92×10^9^.

### 2.7. Karyotyping and genomic abnormalities analysis

Genomic DNA was extracted with the Genomic DNA Mini Kit. Genomic integrity was detected with the hPSC Genetic Analysis Kit (Stemcell, 07550) according to the manufacturer’s instructions. Briefly, 5 ng of genomic DNA was mixed with a ROX reference dye and double-quenched probes tagged with 5-FAM. The probes represented eight common karyotypic abnormalities that have been reported to arise in hiPSC: chr 1q, chr 8q, chr 10p, chr 12p, chr 17q, chr 18q, chr 20q or chr Xp. Sample-probe mixes were analyzed on a QuantStudio 5 Real-Time PCR System (ThermoFisher Scientific). Copy numbers were analyzed using the ΔΔCt method. The results were normalized to the copy number of a control region in chr 4p [20]. For G-band karyotyping, iPSCs were cultured for 72h until they attained 50-60% confluency, then were shipped live to the Wicell Cytogenetics Core (instructions provided by WiCell).

### 2.8. Three Germ Layer Differentiation Test

To form EBs, 2 wells of a 6 well plate containing 80-90% confluent iPSCs were dissociated into small clumps and cultured on low-attachment tissue plates in their preferred iPSC maintenance media (based on CV assays in **2.4**.). On day 7, EBs were transferred to Matrigel-coated plates and left to spontaneously differentiate for 14 days in DMEM media (Wisent, 319-005-CL) containing 20% FBS (Gibco, 10091), 1% NEAA solution (Wisent, 321-011-EL), 0.1 mM 2-mercaptoethanol (Gibco, 31350010) prior to fixation and immunocytochemistry.

To differentiate iPSCs into each of the three germ layers, cells were passaged and dissociated as described above for EB generation into single cells and cultured on Matrigel-coated plates with the STEMdiff Trilineage differentiation kit (used according to the manufacturer’s instructions, Stemcell Technologies, 05230). Cells were harvested at the indicated day for gene expression analysis by qPCR.

### 2.9. Immunocytochemistry analysis

Cells were fixed in 4% PFA/PBS at RT for 20 minutes, permeabilized with 0.2% Triton X-100/PBS for 10 min at RT, then blocked in 5% donkey serum, 1% BSA and 0.05% Triton X-100/ PBS for 2h. Cells were incubated with primary antibodies in blocking buffer overnight at 4 °C. Secondary antibodies were applied for 2h at RT, followed by Hoechst 33342 nucleic acid counterstain for 5 minutes. Immunocytochemistry images were acquired using the automated Evos FL-Auto2 imaging system (ThermoFisher Scientific). Antibodies used for staining are listed in **Table S2**. Images were quantified using custom ImageJ macros. The analysis scripts are available at https://github.com/neuroeddu/CellQ. The thresholds were determined visually by comparing five randomly sampled images.

### 2.10. Cortical neuron differentiation

Differentiation into cortical neurons was based on a protocol for EB formation combined with dual inhibition of SMAD [23, 26] with modifications. Briefly, each iPSC line was cultured in its preferred media for 5-7 days, then dissociated into single cells to form EBs. The EBs were grown in a low-attachment plate for one week in DMEM/F12 supplemented with N2 and B27, in the presence of 10 μM SB431542, and 2μM DMH1. On day 7, EBs were transferred to polyornithine- and laminin-coated plates to form rosettes in the same media. On day 14, rosettes were selected semi–manually and cultured as a monolayer on polyornithine and laminin-coated plates to generate neural progenitor cells (NPCs) in DMEM/F12 supplemented with N2 and B27. NPCs were passaged at a 1:3 dilution every 5-7 days. Immunocytochemistry and qPCR analysis of NPC were conducted at day 25. NPCs were next cultured in neurobasal medium, supplemented with N2 and B27, in the presence of 1 μg/ml laminin, 500 μM db-cAMP, 20 ng/ml BDNF, 20 ng/ml GDNF, 200 μM ascorbic acid, 100 nM Compound A and 1 ng/ml TGF-β for differentiation into neurons. Immunocytochemistry and qPCR analysis of cortical neurons were conducted at day 56.

### 2.11. Data visualization and statistical analysis

All data visualization plots were created in R using the ggplot2 graphical package. For qPCR quantification, the standard deviation (SD) values were calculated in Python. For enhanced visualization of the qPCR quantification in **Figures. 6, S5 and S6** the values were log transformed, and the SD was equivalently scaled. For image analysis SD values were calculated in R. For all plot variation is presented as mean ±SD. A minimum of two independent replicates for each experiment were performed. For calculating the difference in gene expression between the two medias, two-tailed students t-tests were performed using the t.test function in R, individual cell lines were considered biological replicates (n=6 mTeSR1, n=5 E8).

## 3. Results

### 3.1. Validation of hiPSC culture conditions

To work with any iPSC line, it is critical to first establish the optimal growth conditions for that cell-line. As a first step in our workflow, and as part of the initial tests for our newly reprogrammed iPSCs, we grew all our newly generated iPSCs cell-lines side by side in two distinct maintenance media: mTeSR1 or E8. Commercial iPSC lines (NCRM1-NIH and KYOU-DXR0109B-ATCC) were also grown in both media for direct comparison with our “*in house*” control lines (see cell line profiles in **Table 1**). Cells were seeded at an identical confluency and cultured in Matrigel-coated plates in either mTeSR1 or E8 media. We first examined cells 24h after passaging for differences in their overall attachment, spontaneous differentiation and morphology as they grew and expanded. All iPSC lines attached onto Matrigel-coated plates and demonstrated similar morphology when maintained in either media condition. The condensed, round, diffuse and irregularly shape associated with iPSC colonies was observed across all cell lines. The colonies were smooth-edged, with tightly packed cells observed by phase contrast imaging (**Figures. 2A and 3A**). Rare incidences of spontaneous differentiation, in which the cells lost their pluripotency, could be observed with both media conditions and across the different cell-lines.

**Figure 2.**
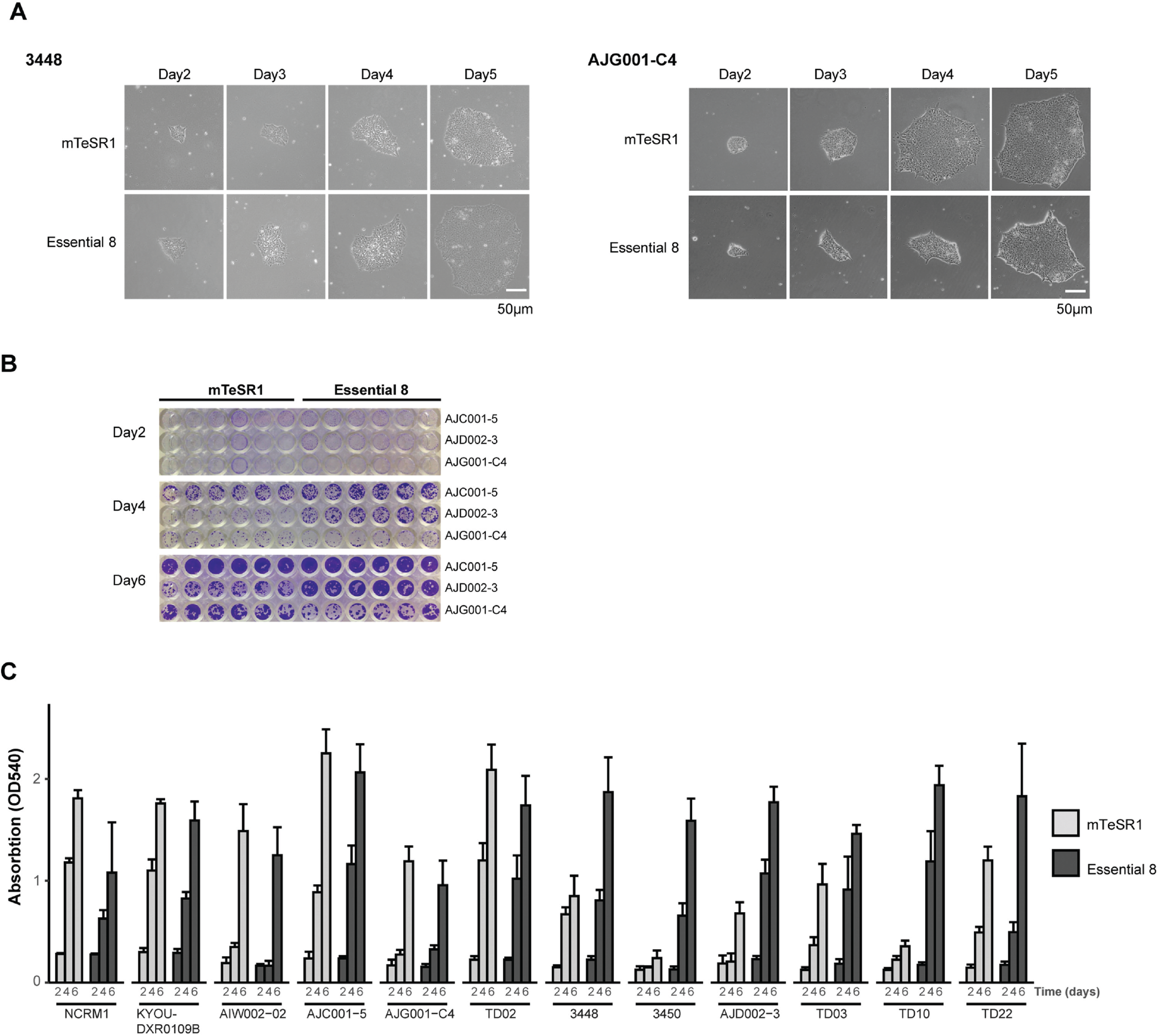
HiPSC growth and proliferation under different conditions. (**A**) Representative daily morphology of hiPSCs maintained in mTeSR1 or E8 media. Both cell lines show smooth-edged, tightly packed cells with a large nucleus-to-cytoplasm ratio. AJG001-C4 cells grows slightly better in mTeSR1 media, while 3448 proliferate better in E8. (**B**) CV assay of representative cell lines grown in mTeSR1 and E8. AJC001-5 grows well in both media. AJG001-C4 grows better in mTeSR1 media, while AJD002-3 cells proliferate best in E8. (**C**) Quantification of the hiPSC’s survival and growth in different media. The CV stain was dissolved in methanol and optical density measured at 540 nm (OD540). The mean and the standard deviation are from 6 replicates from two independent experiments.

**Figure 3.**
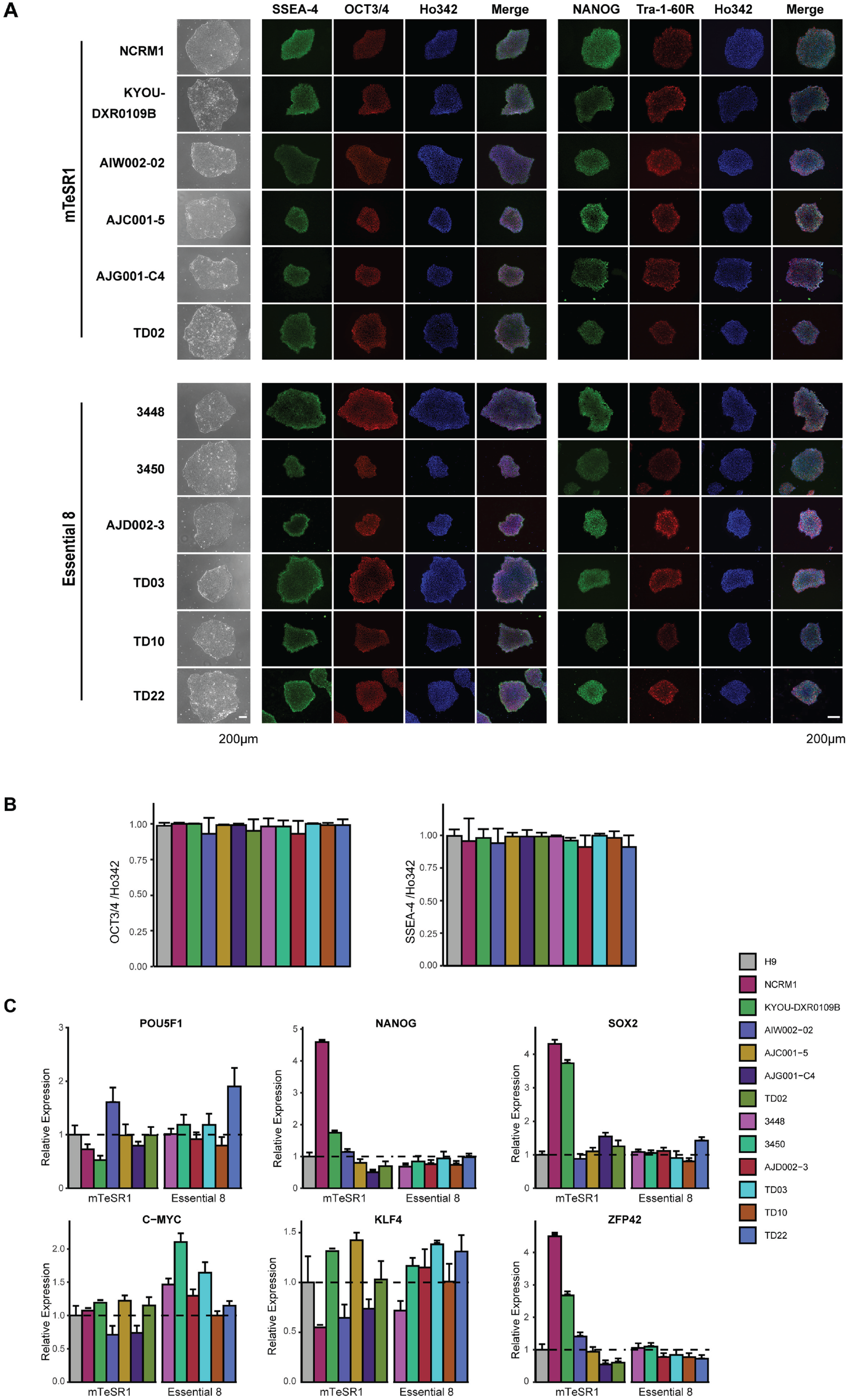
Expression of pluripotency markers in hiPSCs. (**A**) Phase contrast images and ICC for pluripotency markers SSEA-4, OCT3/4, NANOG and Tra-1-60 with Ho342 nucleic acid counterstain. (**B**) Quantification of OCT3/4 positive cells or SSEA-4-positive cells in (**A**). (**C**) qPCR for mRNA expression of pluripotency genes POU5F1, NANOG, c-MYC, KLF4, SOX2 and ZFP42 in hiPSCs compared to H9 ESCs. mRNA expression in H9 was set as 1.0. The mean and SD are from technical triplicates from two independent experiments.

Although morphology is one indicator of iPSC quality, growth properties can vary from line to line, which can be influenced by the media. To detect differences in growth rates and adherence of cells over time, cells were fixed and stained with crystal violet (CV) (**Figures. 2B and S1)** and demonstrate that the growth rate of all cell-lines in each media condition was comparable at day 2. By day 4, three lines were observed to be growing at a faster rate in E8 (AJD002-3, 3450 and TD10; **Figures. 2C and S1**) and exhibited reduced growth and proliferation when maintained in mTeSR1. By day 6, the difference in growth rates was further pronounced for these lines, with 3448 and TD22 also demonstrating a preference for E8. Of note, the proliferation rate for AJG001-C4 was the slowest across the lines, growing at a comparable rate in both media (**Figures. 2C and S1**). For our panel of iPSCs, variations in the growth rate between certain iPSCs was observed depending on the media conditions. Thus, in the rest of our assays, the cells were grown in their preferred media (**Table 1**). Taken together, daily morphological observations coupled with a CV assay, enable a rapid and economical assessment of the growth conditions for a given cell-line, ensuring that each iPSC is cultured in its optimal media for growth and expansion.

### 3.2. Characterization of iPSC pluripotency

Next, we examined the pluripotency of each iPSC line in its preferred growth media (mTeSR1 or E8), as determined by the CV assays (**Figure. 2**). All iPSC lines tested expressed the pluripotency markers SSEA-4, OCT3/4, NANOG and Tra-1-60R, in the same manner as the H9 ESC cell-line as determined by immunohistochemistry (ICC). The cell lines displayed no differences in the fluorescence intensity of these markers (**Figure. 3A**). Notably, we found that over 90% of cells were OCT3/4 or SSEA-4 positive, demonstrating that the cells were maintained in a state of pluripotency with rare spontaneous differentiation (**Figure. 3B**). iPSCs were not only morphologically similar each other and to ESCs but were also similar at a transcriptional level for a number of pluripotency markers that can be commonly observed (OCT3/4, SOX2, NANOG) [27–31]. To examine the transcriptional profile of our lines, the embryonic stem cell, H9 was used as a reference line to normalize the expression of pluripotency genes relative to each of the control iPSC lines [32]. The expression of the Yamanaka factors (OCT3/4, SOX2, KlF4 and c-MYC) and two other widely tested pluripotent markers, NANOG and ZFP42 (Rex1) [33] were analyzed by qPCR (**Figure. 3C**). We found that all iPSC lines expressed each of the pluripotency genes, and expression levels for each gene was comparable to levels in the H9 ESC cell-line. However, variations in expression of the genes was observed between the lines. We found that NCRM1 and KYOU-DXR0109B, two commercial control lines with high passage numbers, expressed relatively high levels of NANOG, SOX2 and ZFP42 when compared to H9, suggesting that prolonged periods of proliferation and self-renewal may stabilize and perpetuate the genome of the hiPSC in a pluripotent state. Given our iPSCs are at an earlier passage number, these levels were lower, consistent with previous studies showing that hESC-specific genes [34] are expressed at lower levels in early passage iPSCs compared to ESCs and late passage iPSC [35].

### 3.3. STR and genomic abnormality testing of hiPSCs

The risk of cell misidentification and cross-contamination has plagued cell research [36, 37]. In addition, hiPSCs which are generated and grown on a layer of mouse embryonic fibroblasts (MEF) feeder cells are often at risk of cross-contamination with non-human rodent somatic cells [38]. For this reason, we conducted STR analysis for iPSC authentication. Our analysis demonstrated that the STR profile for each iPSC tested matched the parental cell-line from which it was reprogrammed, and no rodent contamination was detected with any of our iPSCs. As both AJD002-3 and TD03 were generated from the same donor, they displayed identical STR profiles relative to each other, and to somatic cells obtained from this donor (PBMC sample #3059) (**Table 2**).

**Table 2:**
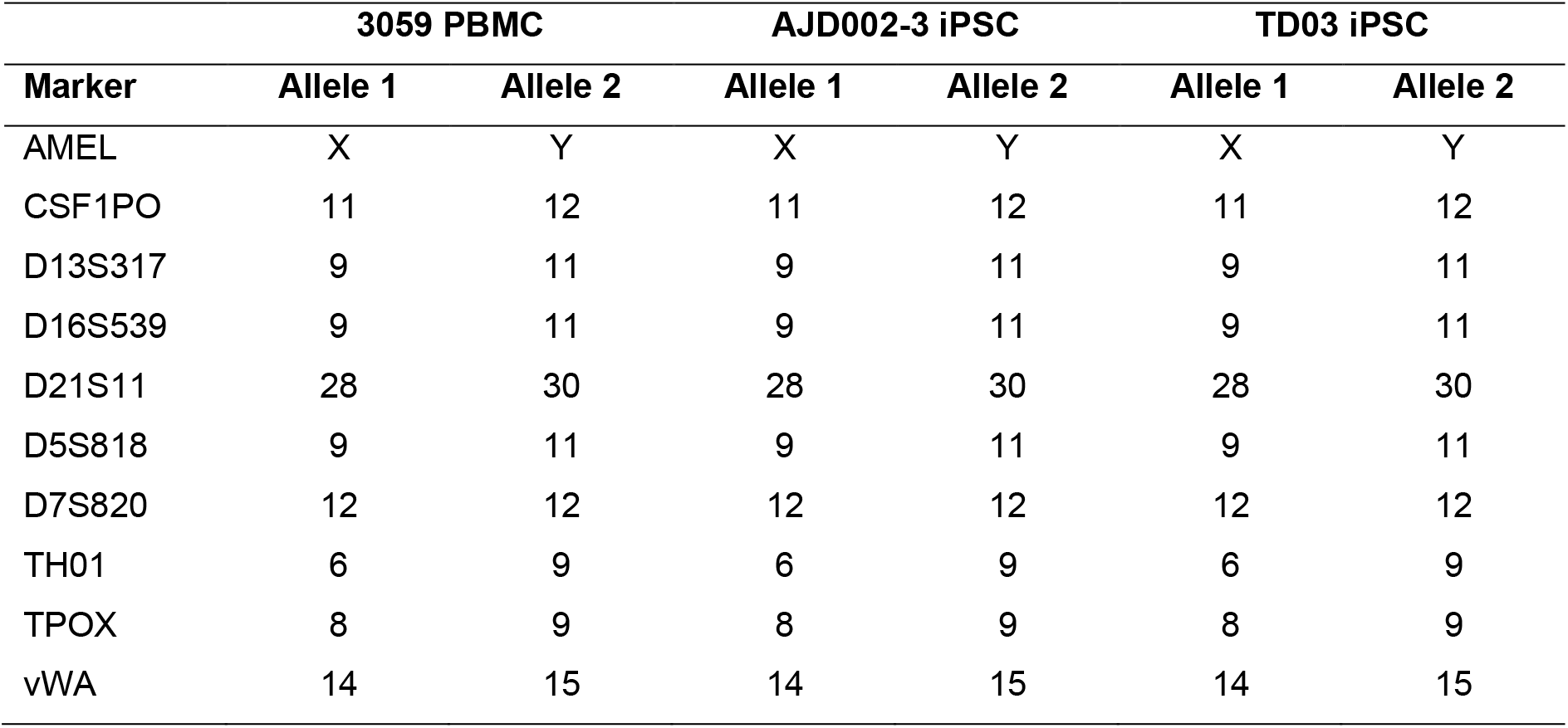
Representative STR profiling of AJD002-3 and TD03 for authentication.

Numerous studies have demonstrated that ESCs and iPSCs accumulate genomic alterations and mutations though reprogramming processes and long-term culture. At the genome level these changes can include copy number variations, trisomy’s, small genomic insertions and deletions, and single nucleotide variants. To assess chromosomal integrity of the iPSC lines, G-band karyotyping was performed. Our analysis showed that the majority of iPSCs tested had a normal 46, XY or 46, XX karyotype (**Figures. 4A and S2**). However, we did detect a chromosomal anomaly in the TD10 line, which contained a translocation between the long (q) arm of chromosome X and the short (p) arm of chromosome 2. These abnormalities were confirmed to be a direct result of reprogramming, as follow-up G-band analysis of the parental PBMCs showed a normal karyotype (data not shown). This confirms that the chromosomal rearrangement in TD10 likely occurred during the reprogramming process and disqualifies this line from further use.

**Figure 4.**
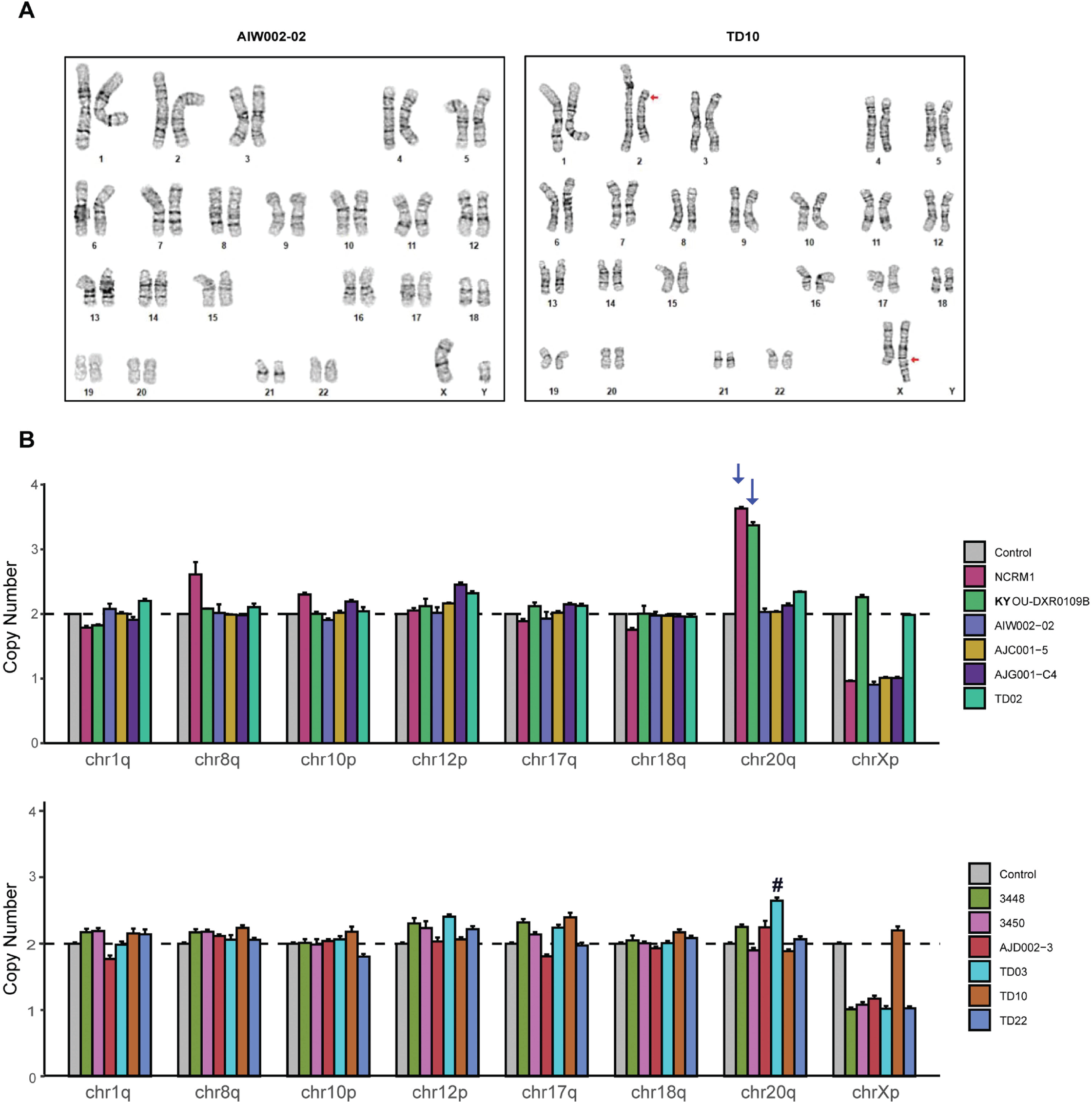
Genomic integrity analysis of hiPSCs. (**A**) Karyotyping and G-band analyses showing examples of one normal (left) and one abnormal hiPSC karyotype (right). An apparently balanced translocation between the long (q) arm of chromosome X and the short (p) arm of chromosome 2 is present in TD10 (red arrow). (**B**) qPCR based genetic analysis to detect small chromosome abnormalities in all hiPSC lines. The copy numbers of chr1q, chr8q, chr10p, chr12p, chr17q, chr18q, chr20q and chrXp were normalized to chr4p expression set to be 2. Error bars show standard deviation from technical triplicates from two independent experiments. There are no abnormalities within critical regions in The Neuro cell lines, while NCRM1 and KYOU-DXR0109 have amplification of chr20q (blue arrow). TD03 shows a slight increase in chr20q, however this is not above the threshold of expected variation.

Although karyotyping by G-banding reveals both numerical and structural aberrations within chromosomes, the limited resolution of this method means we can only detect chromosomal aberrations greater than 5 Mb [14, 39, 40]. To test for commonly occurring genomic alterations, we used a qPCR-based genetic analysis kit to detect minimal critical hotspot regions within the genome that are frequently mutated during the reprogramming process and extended cell passaging, often conferring selective growth advantages to the cells [41, 42]. When testing our newly generated iPSCs, we did not detect any increase or decrease in copy number outside the confidence interval (1.8 to 2.2) indicating there are no abnormalities in any of the eight common hotspot zones tested on chr 1q, chr 8q, chr 10p, chr 12p, chr 17q, chr 18q, chr 20q and chr Xp with our “*in house*” iPSCs, which cover the majority of the reported abnormalities (**Figure. 4B)** [21, 43–46], except for a moderate increase in the copy number for chr 20q in our TD03 line (**Figure. 4B, indicated with #**). However, with both the commercial lines, NCRM1 and KYOU-DXR0109, an amplification in copy number on chromosome 20q was detected (**Figure. 4B, indicated with arrows**), suggesting that this abnormality expanded with extensive cell passage. Thus, it is imperative that when working with any iPSC line, they are profiled for the presence of genomic alterations, which might otherwise affect its growth and differentiation.

### 3.4. Differentiation of hiPSCs into three germ layers

One of the hallmark features of iPSCs is their ability to differentiate into nearly any cell type of the three germ layers when provided with the appropriate developmental cues. To characterize the functional pluripotency of our newly generated iPSCs relative to the commercial lines, we tested their ability to form EBs, in which cells spontaneously differentiate into each of the three embryonic germ layers. After one week, all iPSCs successfully formed EBs and no discernible differences in the relative size and total number of EBs formed was detected between lines. Following the formation of EBs in defined media, they were plated onto Matrigel-coated dishes in the presence of serum-containing media and cultured for 2 weeks. All iPSCs tested could differentiate into each of the three germ layers, as shown by positive immunostaining for the ectoderm marker PAX6, the mesoderm marker SMA and the endoderm marker Vimentin (**Figure. 5A**).

**Figure 5.**
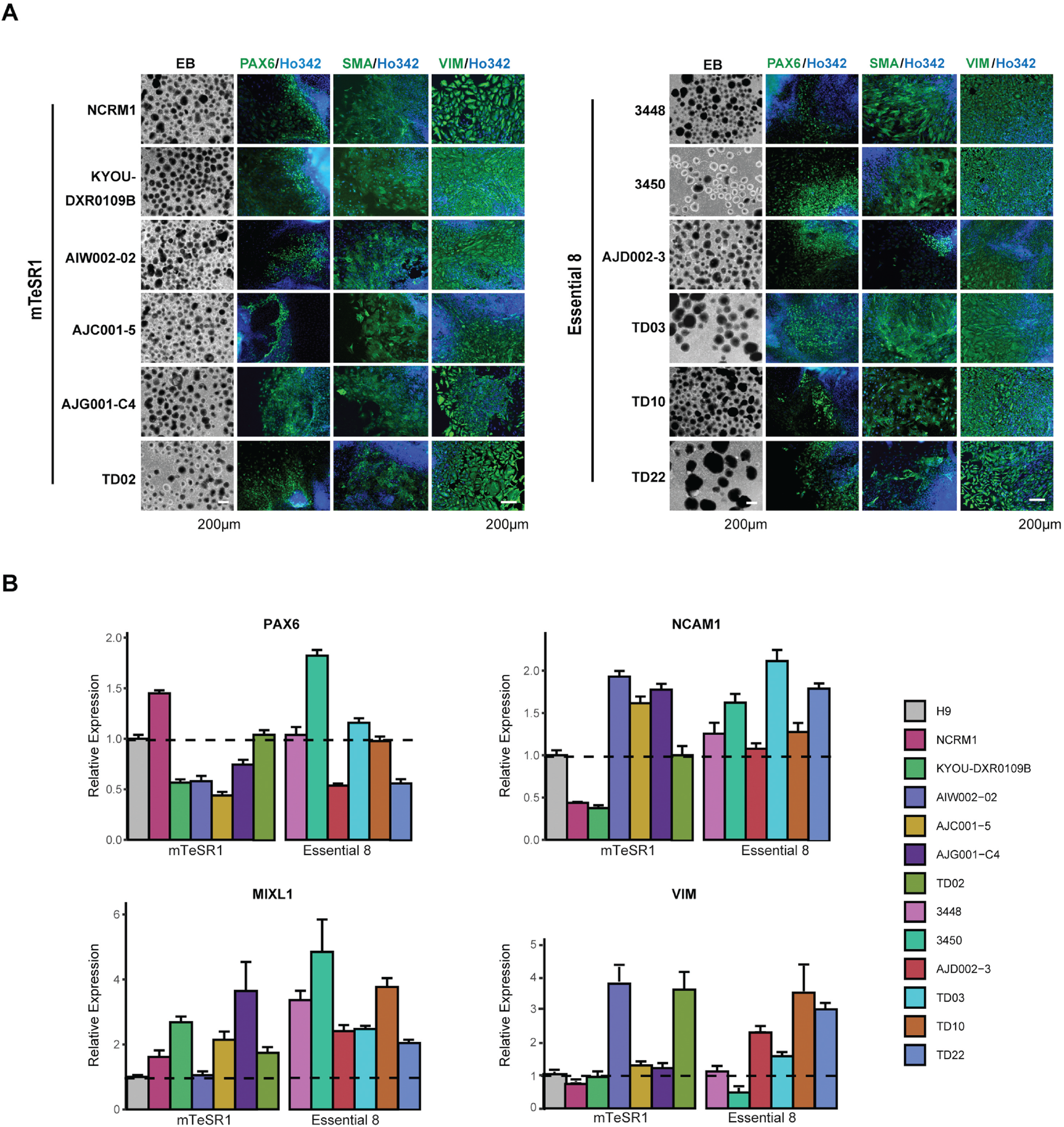
Differentiation of hiPSCs into three germ layers. (**A**) Representative phase contrast images of EB formation (left) and differentiation into three germ layers by ICC with the ectoderm (PAX6), mesoderm (SMA) and endoderm (VIM) markers (indicated above images). Nuclei are counterstained with Ho342. (**B**) qPCR for mRNA expression of three germ layer genes. Quantification of expression of the ectoderm (PAX6, NCAM1), mesoderm (MIXL1) and endoderm (VIM) markers in hiPSCs compared to H9 ESC. mRNA expression in H9 was set as 1.0. The mean and SD are from technical triplicates from two independent experiments.

In parallel to our image analysis, we also took advantage of a qPCR based assay to permit a faster, more quantitative assessment of functional pluripotency. Through this approach, we were able to quantify the *in vitro* differentiation potential of our hiPSCs by measuring the relative expression of key genes that represent each of the three specific lineages. We used the H9 ESC line as our control to compare the expression of three germ layer genes in the iPSC lines. As shown in **Figure. 5B**, cells differentiated from iPSCs into each of the three lineage layers expressed ectodermal (PAX6 and NCAM), mesodermal (MIXL1) or endodermal (Vimentin) markers, depending on their given lineage. However, while all the lines expressed each of these markers, variations were observed between lines, potentially indicating differing capabilities for each line to generate the different germ layers. For instance, 3450 expressed high levels of both Pax6 and MIXL1, while also expressing the lowest levels for Vimentin, indicating this line is likely best used to generate cells of an ectodermal or mesodermal lineage. In contrast, TD10 expressed all the lineage markers at relatively high levels, indicating that the presence of the karyotypic abnormality had no effect on its ability to differentiate and form each of the three germ layers. ICC staining and qPCR findings confirmed that all iPSCs were capable of differentiation into each of the three germ layers. However, based on the expression analysis, the abilities of iPSC to generate different cell types was variable, highlighting the need to choose iPSC lines carefully based on the cell types to be generated for downstream applications.

### 3.5. Differentiation of iPSCs into cortical neurons

Following our trilineage analysis, we narrowed our focus from all germ layers down to one specific cell type from the ectodermal lineage, cortical neurons. To generate cortical neurons, we used a previously published dual SMAD inhibition protocols with modifications [23, 26]. **Figure. 6A** shows the timeline schematic for the protocol, in which iPSCs were differentiated into neurons over 56 days. EBs were formed by dissociating iPSCs at day 0. After 14 days of neural induction, columnar neuroepithelium cells and typical neural tube-like rosettes appeared with all iPSC lines. Rosettes were dissociated and expanded into neural progenitor cells (NPCs). All characterization of NPCs was performed at day 25. We found that all iPSC lines could generate NPCs, as demonstrated by positive immunostaining for the neural progenitor markers Nestin, SOX1, and still maintained an ability to proliferate, as shown by the presence of the cell proliferation marker Ki67 (**Figure. S3**). Based on the quantification of immunostaining, approximately 60-80% of cells from the iPSCs tested were Nestin-positive. In contrast, 10-25% of the cells for each line expressed SOX1 or Ki67 (**Figure. S3B**). Pluripotency (POU1F5 and NANOG), mesodermal (MIXL1) and endodermal (AFP) markers were expressed at low levels, which was to be expected given the cell type generated was of an ectodermal lineage (**Figures. S4A and S4B**). The expression of the NPC marker Nestin was further confirmed by qPCR (**Figure. S4C**). NPCs also expressed dorsal forebrain progenitor (SLC1A3 and PAX6), and ventral forebrain progenitor (ASCL1) markers (**Figure. S4C**), which confirms that the iPSCs tested can differentiate into NPCs [23, 47].

**Figure 6.**
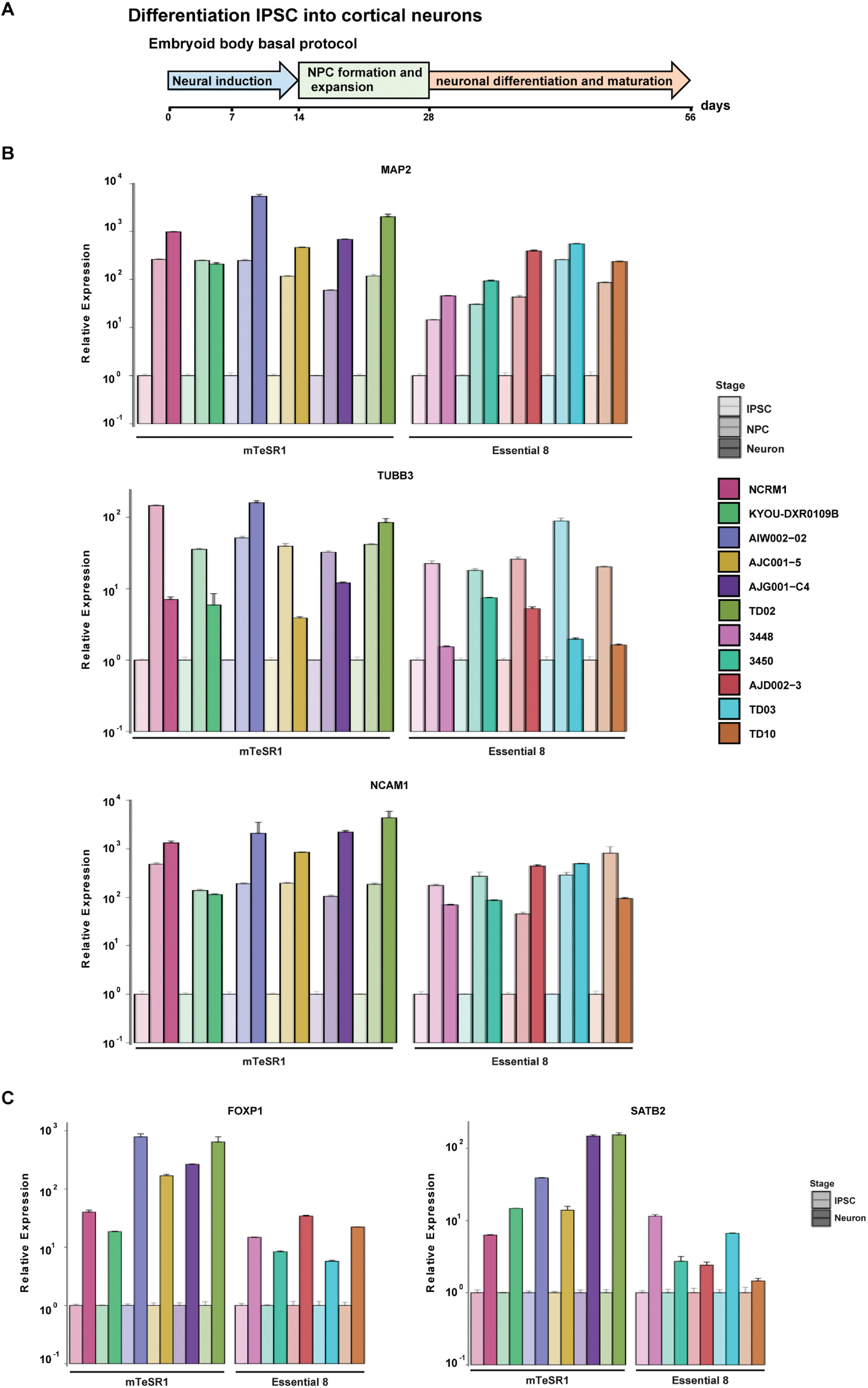
Differentiation of hiPSC into forebrain cortical neurons. (**A**) Schematic representation of the *in vitro* differentiation protocol used to generate cortical neurons from hiPSC. Boxes indicate differentiation state. This protocol was performed in 11 independent lines, with all lines performing similarly. (**B**) qPCR for mRNA expression of neuronal markers MAP2, NCAM1 and TUBB3 in NPC and 4 weeks differentiated cortical neurons compared to IPSCs. mRNA expression in iPSCs was set as 1.0. The mean and SD are from technical triplicates from two independent experiments. (**C**) Expression of cortical layer markers FOXP1 and SATB2 by qPCR analysis of 4 weeks differentiated cortical neurons compared to iPSCs. mRNA expression in iPSCs was set as 1.0. The mean and SD are from technical triplicates from two independent experiments.

NPCs generated from each line were subsequently differentiated into cortical neurons that we analyzed through a combination of qPCR and ICC analysis (**Fig. 6A**). qPCR results confirmed high expression levels of the neuronal markers MAP2, NCAM1 and Tuj (TUBB3) in neurons at day 56 differentiation (**Fig.6B**). Furthermore, upper layer (SATB2) or lower layer (FOXP1) cerebral cortex markers were detected in neurons (**Fig. 6C**). However, the expression level of these markers exhibited line to line variation. Intriguingly, we compared expression levels in the neurons derived from iPSCs lines maintained in mTeSR1 to neurons from iPSCs maintained in E8 and found that levels of these markers was significantly higher for neurons derived from iPSCs maintained in mTeSR1 (p= 0.0349 MAP2; p=0.017 NCAM1; p= 0.018 TUBB3; p = 0.008 FOXP1 and p = 0.010 SATB2) (**Fig. S5**). The higher expression levels of these markers implies that the iPSC maintenance media may influence the differentiation potential of iPSCs, and in this case specifically, into cortical neurons, meriting further investigation. Based on the quantification of the immunostained neurons, approximately 80-90% of cells expressed MAP2 and 60-80% of cells were Tuj1-positive neurons, varying from line to line (**Fig. 7**). In contrast, the expression of the cortical neuron markers Brn2 or Tbr1 was much more variable (5-30%) across cell-lines (**Fig. 7**). Taken together, the iPSCs tested can generate neurons, although variations in morphology and expression of neuronal or cortical region markers does vary across lines. Thus, such variables may similarly impact iPSC differentiation into other cell types making it imperative that before any studies, an iPSC is comprehensively phenotyped to obtain a baseline on its cellular characteristics.

**Figure 7.**
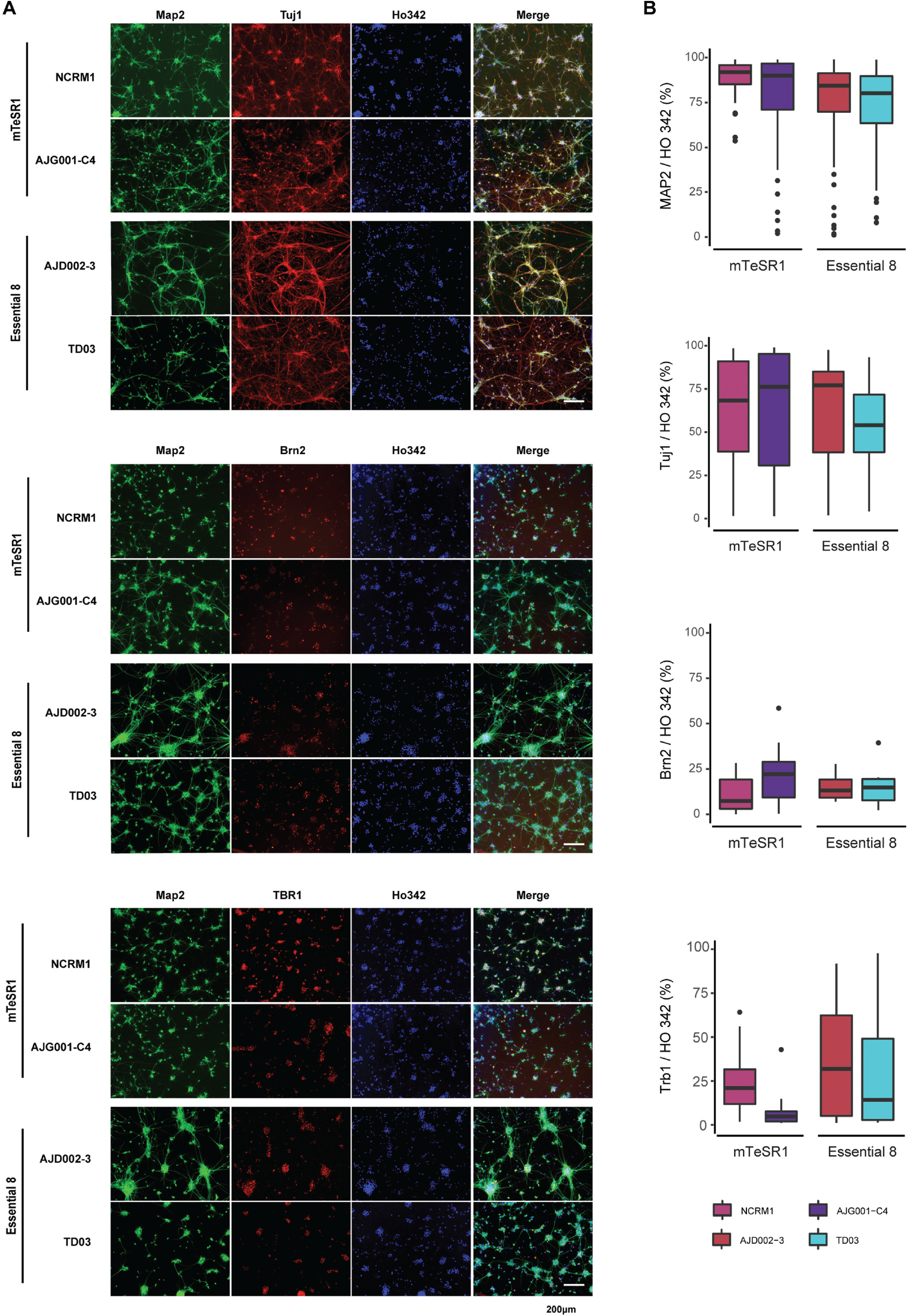
Immunocytochemistry staining of hiPSC-derived cortical neurons. (**A**) Representative images of ICC-stained cortical neurons from AJG001-C4, NCRM1, AJD002-3 and TD03 after 4 weeks neuronal differentiation. Cells were stained with MAP2 and Tuj1 neuronal markers, and cortical neuron markers Brn2 and Tbr1. Nuclei were counterstained with Ho342. (**B**) Quantification of MAP2, Tuj1, Brn2 or Tbr1 positive cells in (**A**).

## 4. Discussion

The routine utilization of iPSCs requires a constant supply of pluripotent, well characterized and quality-controlled cell stocks. However, without standardized quality control, experimental reproducibility with iPSCs can be compromised, making findings difficult to interpret [3, 16, 38]. Here, we established a workflow to monitor the morphology and proliferation of our newly generated iPSCs in two different media (mTeSR1 and E8). In parallel, we evaluated the genomic abnormalities, pluripotency and differentiation potential of our lines. Based on these parameters, we can evaluate whether iPSCs can be used for further applications. However, this is just one iteration of a workflow and the modular nature of the workflow developed means additional quality control tests can be added in further iterations. These can include but are not limited to live cell imaging of iPSCs for growth rate analysis, whole genome sequencing and trilineage analysis through teratoma formation. By focusing on cortical neuron differentiation from the panel of iPSCs, we can also better predict how variable cell-lines are in their ability to generate cortical neurons with the same methods while understanding how the neurons generated might differ from line to line, or from batch to batch within the same line.

Culture conditions can affect the quality, stability and of pluripotency hiPSCs [9, 15, 48, 49]. Today, commercially available medium are widely used to culture hiPSCs [4–8]. We used CV staining [25] which is directly proportional to cell biomass for our assays which provides a affordable and straightforward method to quantify the proliferation of each cell line at different time points. However, other approaches can be performed in addition to, and to complement CV assays, from live cell imaging of iPSC growth rate, to fixed analysis for growth markers through immunocytometry and flow cytometry if desired. While many of our lines grew in both media at comparable rates, some cell lines had a preference for one of the media tested. Five cell lines grew faster in E8 (**Figures. 2B, 2C and S1**), while other lines proliferated at a comparable rate in both media tested. The reason is not clear, but it has been demonstrated previously that the reduced composition of E8 (8 factors) can often elicit a faster growth rate [7]. It is also unclear why both the 3450 and TD10 cell-lines grew well in E8, while their growth appeared to stall in mTeSR1. Interestingly, we found that cell lines derived from the same donor (AJC001-5 and AIW002-02, **Table 1**), albeit different cell types (PBMC vs skin fibroblasts), tended to grow at comparable rates in either media type. Among lines generated from the same donor’s PBMCs (AJD002-3 and TD03), the reprogramming methods (Sendai CytoTune vs. episomal) did not appear to affect the preferred media. Thus, before working with any iPSC it is imperative that culture conditions and media are optimized. Moreover, if a line appears to have a slower growth rate with one media type, it would be recommended to test other media conditions to ensure it’s the growth conditions and not the line itself, accounting for the poor growth rate of a given iPSC line.

Expression of pluripotency-associated markers is an important quality criterion for any iPSC line, otherwise the iPSCs cannot be differentiated into a cell type of interest. All iPSC lines in our study expressed the pluripotency markers SSEA-4, OCT3/4, NANOG and TRA-1-60R in defined culture conditions (**Figures. 3A and B**). We did find that OCT3/4 expression level was approximately 15 times higher than that of SOX2 (**Table S3**), consistent with previous studies, showing this OCT3/4 high, SOX2 low stoichiometry is important not only in the early phase of reaching a fully reprogrammed state, but also in the late phase of iPSC maturation and maintaining pluripotency [50–52]. Further evidence has also demonstrated that OCT3/4 downregulates the downstream gene expression of NANOG, SPP1/ OPN, SOX2, FBXO15, OTX2, and ZFP42/REX1 [53]. The expression levels of NANOG, ZFP42 and c-MYC in our study were approximately 50 times lower than that of OCT3/4, which is consistent with previous results from single cell RNA-sequencing iPSC dataset [54, 55]. Interestingly, NCRM1 and KYOU-DXR0109B, two commercial control lines with high passage number (>30 passages), expressed relatively high levels of NANOG, SOX2 and ZFP42 and reduced levels of OCT3/4, suggesting that extended passaging enhances pluripotent gene expression in an undifferentiated state and increases the efficiency of neuronal conversion [56].

A core feature of hiPSCs is their pluripotency, that is, the ability to differentiate into nearly any cell type of the three germ layers. The previous gold standard method to assess pluripotency of iPSCs was a teratoma assay, in which iPSCs were injected into immune-deficient mice to assess their ability to form teratomas [27, 57]. However, this requires the sacrifice of animals, and can be expensive and time consuming, leading to the development of trilineage assays to assess the pluripotency of the cells [58]. One such approach is the *In vitro* formation of EBs, a commonly used method to assess the differentiation capability of a given iPSC [59, 60]. All the iPSCs we tested in our panel had the potential to differentiate into each of the three germ layers, as shown by positive immunostaining for the ectoderm, mesoderm, and endoderm markers PAX6, SMA and Vimentin, respectively (**Figure. 5A**), as previously reported [61]. However, while we successfully generated each of the three germ layers, the heterogeneous nature of the EBs resulted in inefficient and often variable differentiation of the three germ layers with each line. To further standardize our protocols, we used a commercial trilineage differentiation kit to perform parallel *in vitro* directed differentiation experiments for each germ layer. We also took advantage of a qPCR-based assay to enable a faster, more quantitative assessment of functional pluripotency, relative to the image-based approach of the EB trilineage test. Through this approach, we could quantify the *in vitro* differentiation potential of our iPSCs by measuring the relative expression of key genes that represent each of the three lineages. From this test, all the iPSCs expressed markers for each of the three germ layers, albeit at differing levels, with some lines expressing higher levels of one or more markers compared to each other (**Figure. 5B**). Nevertheless, depending on the line itself, some consideration needs to be given as to whether it is the optimal line required to generate a particular cell type of interest.

Numerous studies have demonstrated that both ESCs and iPSCs accumulate genomic abnormalities during long-term culturing, and often is the primary reason why clinical therapies from stem cells were not administered to patients [62]. The presence of genetic variations in iPSCs has raised serious safety concerns for both patient interventions and basic research studies, hampering the advancement of novel iPSC-based therapies. G-banding was widely used for genetic evaluation [14] and upon karyotyping, the majority of iPSCs we tested maintained a normal 46, XY or 46, XX karyotype (**Figures. 4A and S2**). However, we were able to detect a genomic anomaly in one of our control lines, TD10 which presented with an abnormal karyotype, that is, a translocation between the long (q) arm of chromosome X and the short (p) arm of chromosome 2. Yet, when compared to other iPSCs, this line appears within the normal range for other parameters, highlighting how important it is to assess each line for genomic abnormalities with multiple tests. One such test we applied in our analysis was a qPCR-based genetic analysis kit to detect minimal critical hotspot regions within the genome that can arise during the reprogramming process or confer selective growth advantages for a given cell-line [41, 42]. Using this analysis, which covers the majority of reported abnormalities, we did not detect any abnormalities in any of the hotspot zones with our newly generated iPSCs (**Figure. 4B**) [21, 43–46]. However, with the two commercial lines, NCRM1 and KYOU-DXR0109, an amplification in copy number on chromosome 20q was detected (**Figure. 4B, labeled with an arrow**). This region is enriched with genes associated with pluripotency and anti-apoptosis, including DNA methyl-transferase 3B (DNMT3B), inhibitor of DNA binding 1 (ID1), and BCL2-like1 (BCL2L1) [45, 63]. We also detected a slight increase in copy number for chr8q in NCRM1, which is a previously reported abnormality acquired during prolonged periods in culture [14], suggesting rigorous quality control is needed with these lines, as it is likely that with increasing cell passage, these abnormalities have expanded. These findings strongly suggest that it is critical to test lines for genomic abnormalities that can arise through reprogramming and prolonged cell passage. Additional quality control tests worth pursued in future iPSC profiling is whole genome sequencing (WGS) [64, 65]. With recent advances significantly reducing the cost of WGS to more affordable levels for its widespread use in research labs, it can help in future workflows to detect low frequency variations which could not be identified by conventional methods and adding an extra layer of quality control profiling.

Moving beyond a broad trilineage test, we next tested our lines for their ability to form one specific cell type of interest, that is, cortical neurons. Prior studies have shown that dual SMAD inhibitors synergistically destabilize the activin- and NANOG-mediated pluripotency network [66], suppresses BMP-induced mes-/endodermal fates differentiation [67, 68] and promoted neuralization of the primitive ectoderm by BMP inhibition [23]. Taken together, our findings indicate no preference for a specific layer in the generation of cortical neuron by dual SMAD inhibition EB method, yet the expression levels were highly variable across iPSCs. The variability does not come from the differentiation protocol, as all the iPSCs were differentiated under the same conditions. This variability may not simply be a direct result of distinct genetic background differences, since variations in differentiation were also detected between AIW002-02, AJC001-5 and AJG001-C4 which share the same genetic background. Intriguingly, the iPSCs maintained in mTeSR1, expressed FOXP1 and SATB2 at levels much higher than those maintained in E8, potentially indicating that the conditions we culture the cells might impact their ability to form defined types of neurons, an area which warrants further study.

## 5. Conclusions

Reprogramming of somatic cells into iPSCs opens up the possibility of modeling human diseases and developing new therapeutics. Using human iPSCs-derived cells for preclinical and clinical research will require a constant supply of well characterized pluripotent cell-lines. Thus, in this study, we established a workflow to monitor the growth and morphology of newly generated iPSCs in two different media. We also performed a comprehensive phenotyping of the iPSC lines through growth rate profiling, testing of genome integrity, analysis of pluripotency capacity and tests on each of the iPSCs to form each of the three germ layers, with a particular focus on cortical neurons of the ectodermal lineage. From these studies, we demonstrated that our newly generated iPSC lines share common hallmarks yet can vary in their growth rate or ability to differentiate into other cell types. Given our findings, it is imperative that each new iPSC line be evaluated thoroughly before using it in downstream applications, while ensuring the line can be used to generate the cell types of interest for a given research application. With these parameters in mind, the workflow outlined will help streamline work processes and offers the potential to add new tests as technologies evolve, to ensure researchers employ iPSCs of the highest quality for experimental reproductivity and robustness.

## Abbreviations

5-FAM: 5-carboxyfluorescein
CV: crystal violet
E8: Essential 8 media
EB: embryoid body
ESC: embryonic stem cell
iPSC: induced pluripotent stem cell
Ho342: Hoechst 33342
ICC: immunocytochemistry
NE: neuroepithelial
NEAA: non-essential amino acids
NPC: neural progenitor cell
PBMC: peripheral blood mononuclear cells
PFA: paraformaldehyde
RT: room temperature

## Supplementary Materials

**Figure S1**. HiPSC growth and proliferation profile in mTeSR1 or Essential 8 media; **Figure S2**. HiPSCs maintain a normal karyotype; **Figure S3**. Characterization of hiPSC-derived NPCs; **Figure S4**. Gene expression profiles in neural progenitor cells; **Figure S5**. Comparison of gene expression levels in the neurons derived from iPSCs lines maintained in mTeSR1 to neurons from iPSCs maintained in E8; **Table S1**. Primers used in qPCR experiments; **Table S2**. Key resources; **Table S3**. Quantification of pluripotency gene expression in hiPSCs by qPCR analysis.

## Author Contributions

Author contributions were diverse and covered many aspects of the work performed throughout this manuscript and have been broken down following the CRediT taxonomy as follows: Conceptualization, C.C., T.D.; Methodology, C.C., T.D.; Software G.M., R.T., E.C., I.V.; Validation, C.C., N.A., R.T., T.D.; Formal analysis, C.C., N.A., R.T.; Investigation, C.C., N.A.; Resources, J.K., M.T.; Data curation, C.C., N.A., R.T., G. M., E.C.; Writing—original draft, C.C., N.A., L.B.; Writing—review and editing, C.C., T.D., L.B., R.T., E.F.; Visualization, C.C., T.D., L. B.; Supervision, T.D., E.F.; Project Administration, C.C., L.B., T.D.; Funding acquisition, T.D., E.F.; All authors have read and agreed to the published version of the manuscript.

## Institutional Review Board Statement

All subjects gave their informed consent for inclusion before they participated in the study. The study was conducted in accordance with the Declaration of Helsinki, and the protocol was approved by the McGill University Health Centre Research Ethics Board (DURCAN_IPSC / 2019-5374).

## Informed Consent Statement

Informed consent was obtained from all subjects involved in the study.

## Data Availability Statement

Data are not available for distribution from this study.

## Acknowledgments

We thank Drs. Mark Aurousseau and Wolfgang Reintsch for automated image acquisition, Angela Nauleau-Javaudin for RNA sample preparation and Dr. Chanshuai Han for technical advice. Mention of trade names or commercial products in this publication is solely for the purpose of providing specific information and does not imply recommendation or endorsement.

---

**Table S1.** Primers used in qPCR experiments.

| **Gene ID** | **Gene Name** | **Accession number** | **Assay ID** |
| --- | --- | --- | --- |
| **ACTB** | Actin, beta | NM_001101.3 | Hs01060665_g1 |
| **AFP** | Alpha fetoprotein | NM_001134.2 | Hs01040598_m1 |
| **ASCL1** | Achaete-scute family bHLH transcription factor 1 | NM_004316.3 | Hs00269932_m1 |
| **MYC** | v-myc avian myelocytomatosis viral oncogene homolog | NM_002467.4 | Hs00153408_m1 |
| **GAPDH** | Glyceraldehyde-3-phosphate dehydrogenase | NM_001256788.2 | Hs02786624_g1 |
| **KLF4** | Kruppel like factor 4 | NM_001314052.1 | Hs00358836_m1 |
| **MAP2** | Microtubule associated protein 2 | NM_001039538.1 | Hs00258900_m1 |
| **MIXL1** | Mix paired-like homeobox | NM_031944.2 | Hs00430824_g1 |
| **NANOG** | NANOG homeobox | NM_001297698.1 | Hs02387400_g1 |
| **NCAM1** | Neural cell adhesion molecule 1 | NM_000615.6 | Hs00941830_m1 |
| **NES** | Nestin | NM_006617.1 | Hs04187831_g1 |
| **PAX6** | Paired box 6 | NM_000280.4 | Hs01088114_m1 |
| **POU5F1 (OCT3/4)** | POU class 5 homeobox 1 | NM_001173531.2 | Hs04260367_gH |
| **SLC1A3** | Solute carrier family 1 member 3 | NM_001166695.2 | Hs00904823_g1 |
| **SOX2** | SRY-box 2 | NM_003106.3 | Hs04234836_s1 |
| **TUBB3** | Tubulin beta 3 class III | NM_001197181.1 | Hs00801390_s1 |
| **VIM** | Vimentin | NM_003380.3 | Hs00958111_m1 |
| **ZFP42** | ZFP42 zinc finger protein | NM_001304358.1 | Hs00399279_m1 |

**Table S2.** Overview of Antibodies.

| **Antibody** | **SOURCE** | **IDENTIFIER** |
| --- | --- | --- |
| **Rabbit polyclonal anti-Brn2/POU3F2** | Cell Signaling | Cat#12137; RRID: AB_2797827 |
| **Mouse monoclonal anti-CD44** | BD Biosciences | Cat#550392; RRID: AB_2074674 |
| **Mouse monoclonal anti-Ki-67** | BD Biosciences | Cat#556003; RRID: AB_396287 |
| **Chicken polyclonal anti-MAP2** | EnCor Biotechnology | Cat#CPCA-MAP2; RRID: AB_2138173 |
| **Rabbit polyclonal anti-Nanog** | Abcam | Cat#ab21624; RRID: AB_446437 |
| **Rabbit polyclonal anti-Nestin** | Abcam | Cat#ab92391; RRID: AB_10561437 |
| **Goat polyclonal anti-Oct-3/4(N-19)** | Santa Cruz Biotechnology | Cat#sc-8628; RRID: AB_653551 |
| **Mouse monoclonal anti-PAX6** | DSHB | Cat#AB_528427; RRID: AB_528427 |
| **Rabbit polyclonal anti-SMA1** | Abcam | Cat#ab5694; RRID: AB_2223021 |
| **Goat polyclonal anti-SOX1** | R&D Systems | Cat#AF3369; RRID: AB_2239879 |
| **Rabbit polyclonal anti-SOX9** | Millipore | Cat#AB5535; RRID: AB_2239761 |
| **Mouse monoclonal anti-SSEA-4(813-70)** | Santa Cruz Biotechnology | Cat#sc-21704; RRID: AB_628289 |
| **Rabbit monoclonal anti-TBR1** | Abcam | Cat#ab31940; RRID: AB_2200219 |
| **Mouse monoclonal anti-TRA-1-60R** | Stemcell Technologies | Cat#60064; RRID: AB_2686905 |
| **Rabbit monoclonal anti-Vimentin** | Abcam | Cat#ab92547; RRID: AB_10562134 |
| **Mouse monoclonal anti-Tubulin- βIII** | Millipore | Cat#MAB5564; RRID: AB_11212768 |
| **Hoechst 33342** | ThermoFisher Scientific | H3570 |

**Table S3.** Quantification of pluripotency gene expression in hiPSCs by qPCR analysis (mean ± SEM of relative gene expression, normalized to GAPDH and ACTIN)

| Cell line | OCT3/4 | SOX2 | NANOG | c-MYC | ZFP42 | KLF4 |
| --- | --- | --- | --- | --- | --- | --- |
| H9 | 2.219±0.310 | 0.1400±0.0065 | 0.0657±0.0044 | 0.0420±0.0044 | 0.0524±0.0048 | 0.000157±0.000037 |
| NCRM1 | 1.614±0.135 | 0.6026±0.0105 | 0.3019±0.0012 | 0.0661±0.0034 | 0.2360±0.0015 | 0.000042±0.000009 |
| KYOU-DXR0109B | 1.168±0.108 | 0.5223±0.0061 | 0.1148±0.0008 | 0.0998±0.0013 | 0.1403±0.0023 | 0.000082±0.000014 |
| AIW002-02 | 3.567±0.298 | 0.1226±0.0129 | 0.0748±0.0025 | 0.0298±0.0040 | 0.0737±0.0027 | 0.000101±0.000017 |
| AJC001-5 | 2.194±0.383 | 0.1540±0.0081 | 0.0525±0.0036 | 0.0513±0.0017 | 0.0491±0.0035 | 0.000224±0.000008 |
| AJG001-C4 | 1.763±0.096 | 0.2163±0.0087 | 0.0332±0.0016 | 0.0310±0.0029 | 0.0287±0.0024 | 0.000116±0.000011 |
| TD02 | 2.199±0.263 | 0.1753±0.0170 | 0.0458±0.0062 | 0.0483±0.0036 | 0.0316±0.0024 | 0.000162±0.000025 |
| 3448 | 2.248±0.144 | 0.1515±0.0045 | 0.0446±0.0014 | 0.0615±0.0024 | 0.0552±0.0041 | 0.000113±0.000011 |
| 3450 | 2.633±0.345 | 0.1475±0.0053 | 0.0554±0.0061 | 0.0883±0.0042 | 0.0574±0.0026 | 0.000183±0.000009 |
| AJD002-3 | 2.036±0.209 | 0.1556±0.0080 | 0.0499±0.0032 | 0.0545±0.0027 | 0.0405±0.0028 | 0.000181±0.000025 |
| TD03 | 2.626±0.406 | 0.1267±0.0208 | 0.0616±0.0089 | 0.0689±0.0054 | 0.0440±0.0050 | 0.000217±0.000002 |
| TD10 | 1.771±0.274 | 0.1128±0.0057 | 0.0484±0.0025 | 0.0420±0.0015 | 0.0404±0.0032 | 0.000158±0.000024 |
| TD22 | 4.221±0.688 | 0.1991±0.0066 | 0.0641±0.0023 | 0.0482±0.0016 | 0.0378±0.0027 | 0.000206±0.000022 |

## Supplementary Figure Legends

**Figure S1.**
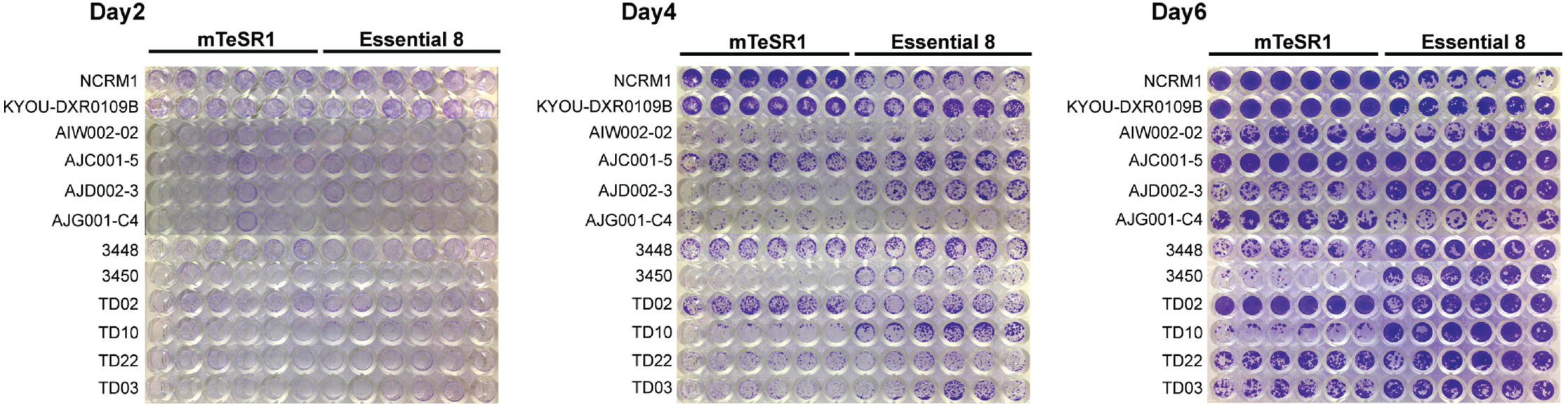
HiPSC growth and proliferation profile in mTeSR1 or Essential 8 media. Related to Fig. 1. Crystal violet staining of hiPSCs grown in mTeSR1 and Essential 8 after day 2, 4 and 6 in culture.

**Figure S2.**
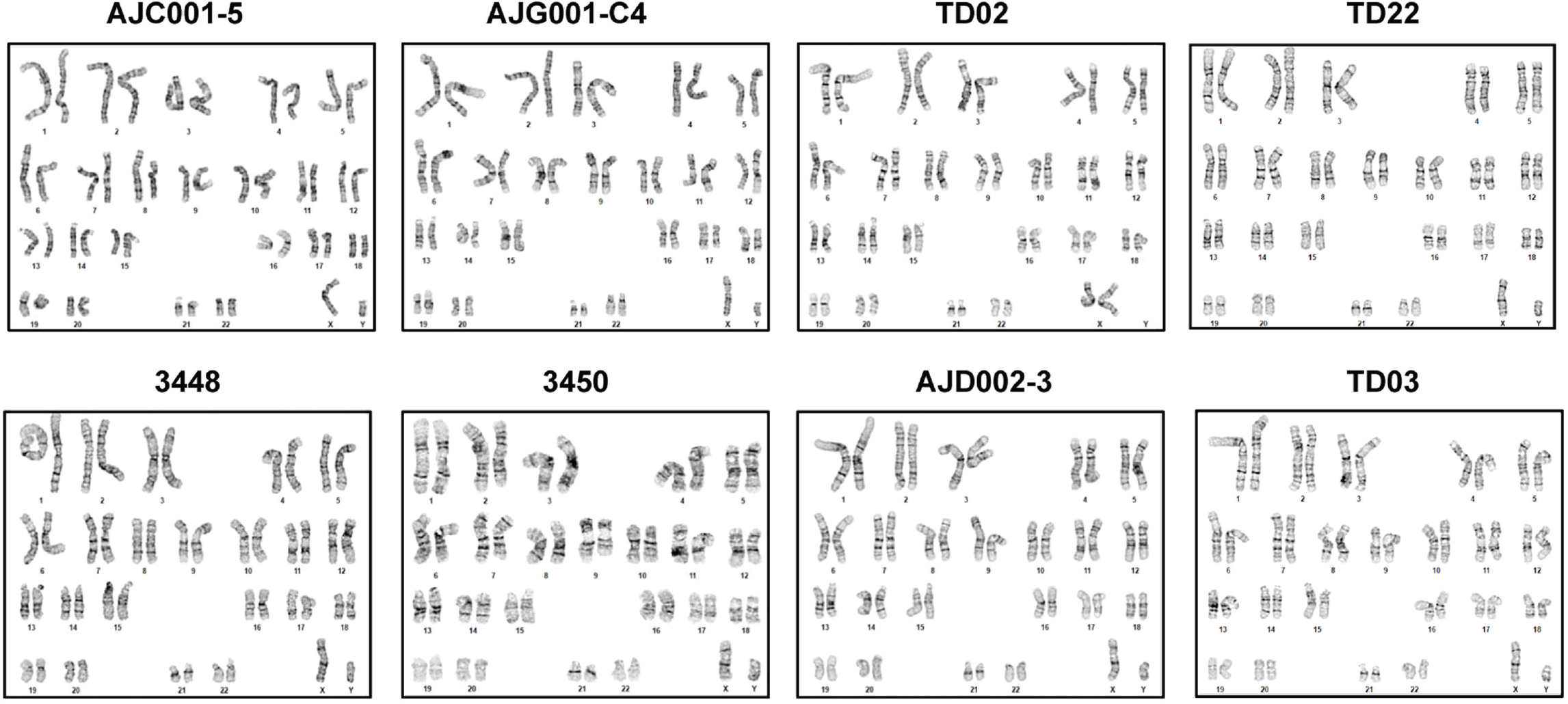
HiPSCs maintain a normal karyotype. Related to Fig. 3. Karyotyping and G-band analyses show 8 control hiPSC lines have normal 46, XY or 46, XX karyotypes.

**Figure S3.**
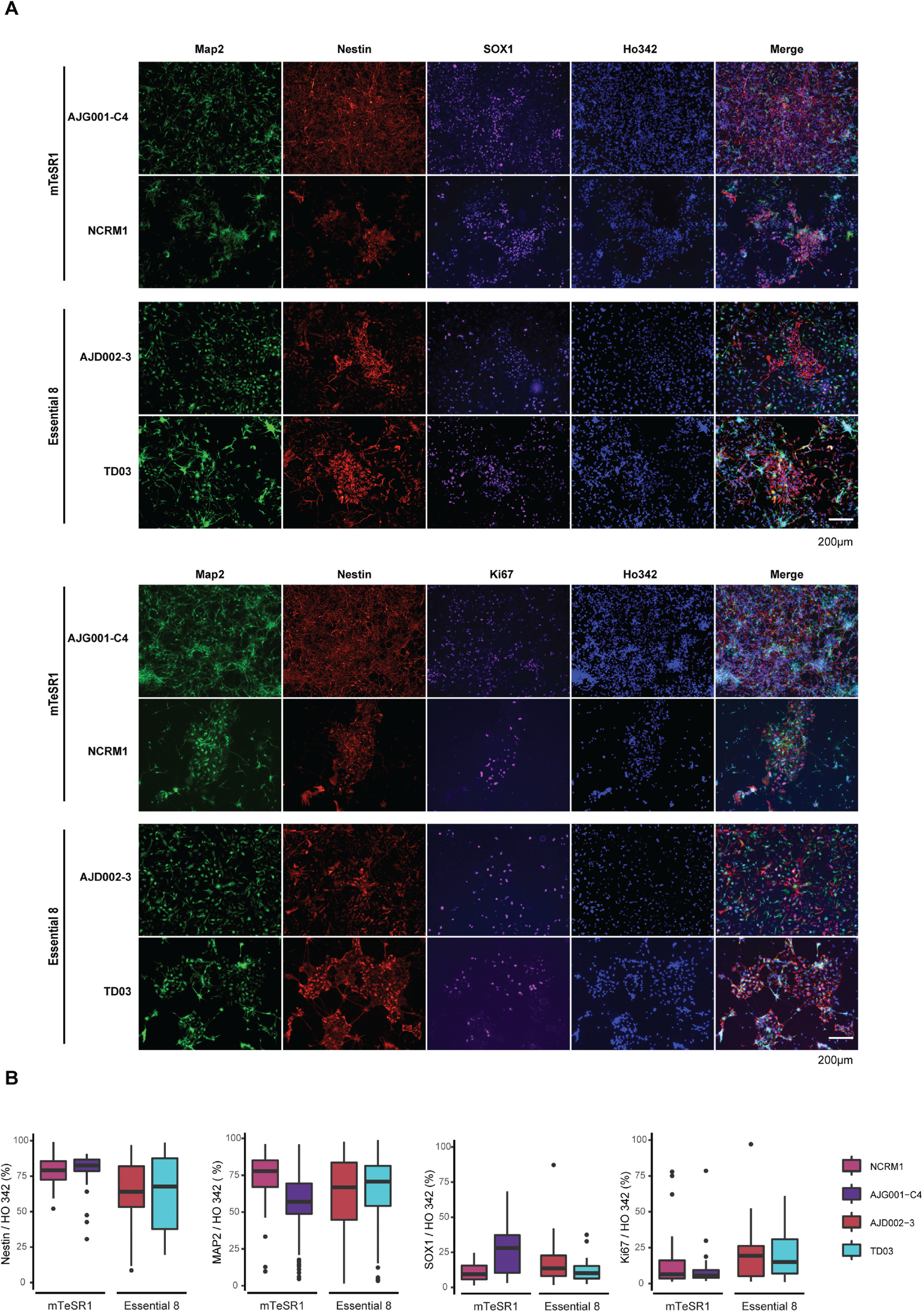
Characterization of hiPSC-derived NPCs. (**A**) Representative images of immunostained NPCs from AJG001-C4, NCRM1, AJD002-3 and TD03 hiPSC. Cells are stained with neural progenitor markers Nestin and SOX1, proliferative marker Ki67 and neuronal marker MAP2. Nuclei were counterstained with Ho342 at day 25. (**B**) Quantification of Nestin, MAP2, SOX1 or Ki67 positive cells in (**B**).

**Figure S4.**
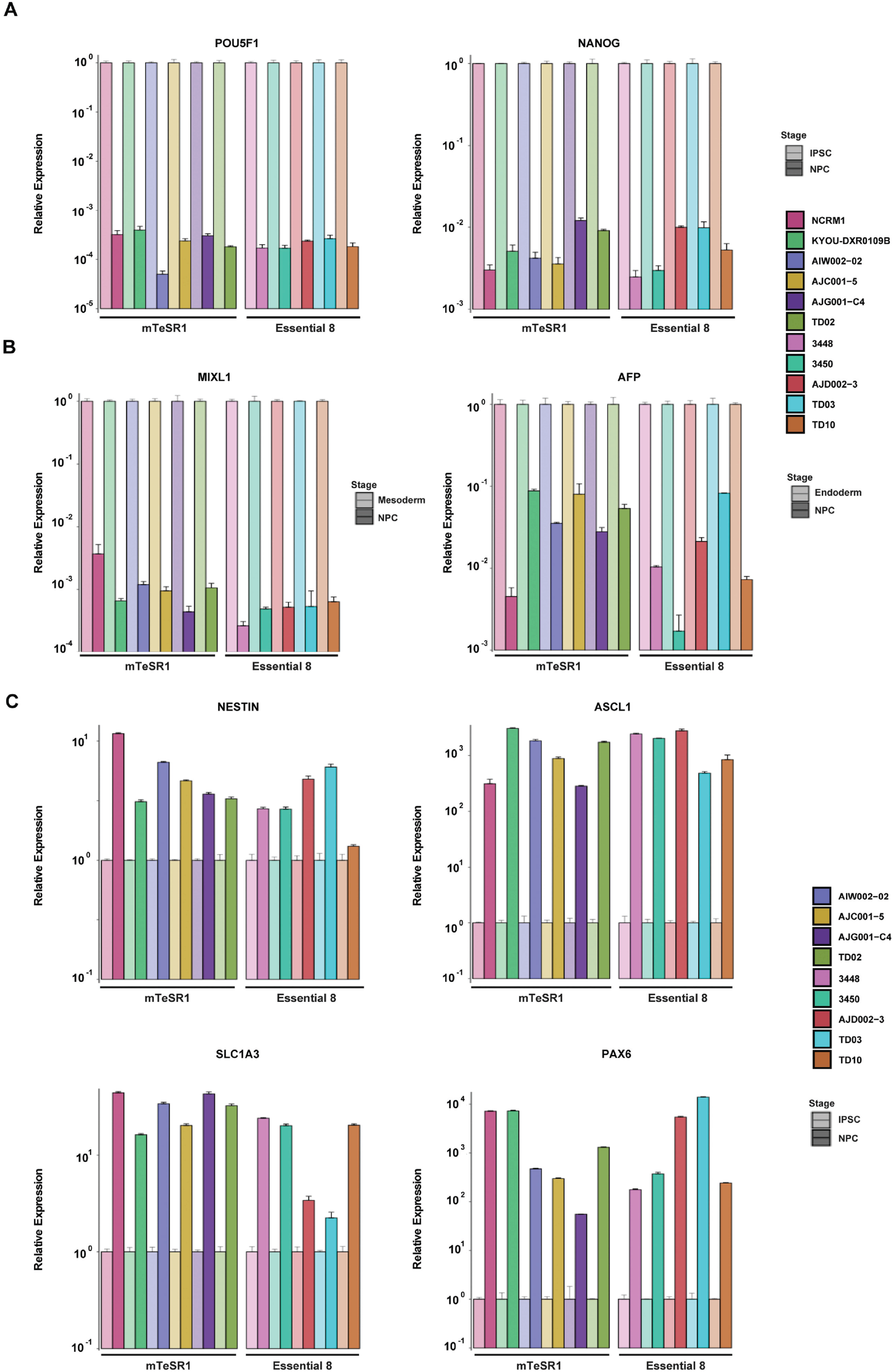
Gene expression profiles in neural progenitor cells. (**A**) qPCR for the mRNA expression of pluripotency markers POU5F1 and NANOG in NPC at day 25 of differentiation. The mRNA expression in iPSCs was set as 1.0. The mean and SD are from technical triplicates from two independent experiments. (**B**) qPCR for the mRNA expression of mesoderm marker MIXL1 and endoderm marker AFP in NPC at day 25 of differentiation. mRNA expression in mesoderm or endoderm was set as 1.0. The mean and SD are from technical triplicates from two independent experiments. (**B**) Quantification of expression of neural progenitor maker Nestin, and the forebrain cortical NPC markers SLC1A3, PAX6 and ASCL1 as measured by qPCR in NPC are compared to iPSCs. mRNA expression in iPSCs was set as 1.0. The mean and SD are from technical triplicates from two independent experiments.

**Figure S5.**
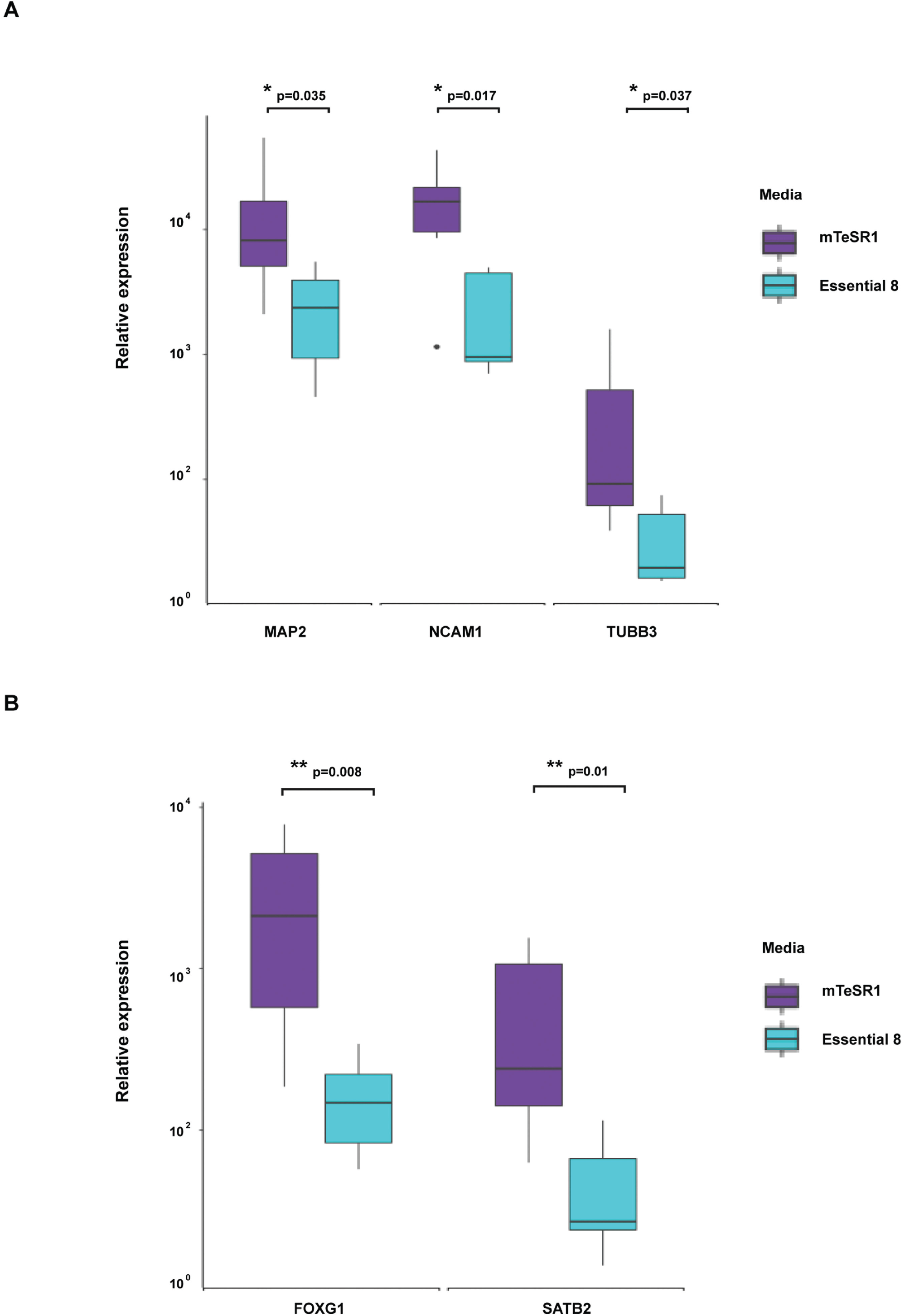
Comparison of gene expression levels in the neurons derived from iPSCs lines maintained in mTeSR1 to neurons from iPSCs maintained in E8. (**A**) The mRNA expression levels of neuronal markers MAP2, NCAM1 and TUBB3 are significantly higher in the neurons derived from iPSCs lines maintained in mTeSR1 (*: p<0.05). (**B**) The mRNA expression levels of cortical layers markers FOXG1 and SATB2 are significantly higher in the neurons derived from iPSCs lines maintained in mTeSR1 than that in Essential 8 (**: p<0.01).

## References

1. Daley GQ. The promise and perils of stem cell therapeutics. Cell Stem Cell. 2012;10(6):740–9. Epub 2012/06/19. doi: 10.1016/j.stem.2012.05.010. PubMed PMID: 22704514; PubMed Central PMCID: PMCPMC3629702.

2. Dolgin E. Putting stem cells to the test. Nat Med. 2010;16(12):1354–7. Epub 2010/12/08. doi: 10.1038/nm1210-1354. PubMed PMID: 21135828.

3. Sullivan S, Stacey GN, Akazawa C, Aoyama N, Baptista R, Bedford P, et al. Quality control guidelines for clinical-grade human induced pluripotent stem cell lines. Regen Med. 2018;13(7):859–66. Epub 2018/09/13. doi: 10.2217/rme-2018-0095. PubMed PMID: 30205750.

4. Ludwig TE, Levenstein ME, Jones JM, Berggren WT, Mitchen ER, Frane JL, et al. Derivation of human embryonic stem cells in defined conditions. Nat Biotechnol. 2006;24(2):185–7. Epub 2006/01/03. doi: 10.1038/nbt1177. PubMed PMID: 16388305.

5. Chen G, Gulbranson DR, Hou Z, Bolin JM, Ruotti V, Probasco MD, et al. Chemically defined conditions for human iPSC derivation and culture. Nat Methods. 2011;8(5):424–9. Epub 2011/04/12. doi: 10.1038/nmeth.1593. PubMed PMID: 21478862; PubMed Central PMCID: PMCPMC3084903.

6. Hey CAB, Saltokova KB, Bisgaard HC, Moller LB. Comparison of two different culture conditions for derivation of early hiPSC. Cell Biol Int. 2018;42(11):1467–73. Epub 2018/04/01. doi: 10.1002/cbin.10966. PubMed PMID: 29603519.

7. Tano K, Yasuda S, Kuroda T, Saito H, Umezawa A, Sato Y. A novel in vitro method for detecting undifferentiated human pluripotent stem cells as impurities in cell therapy products using a highly efficient culture system. PLoS One. 2014;9(10):e110496. Epub 2014/10/28. doi: 10.1371/journal.pone.0110496. PubMed PMID: 25347300; PubMed Central PMCID: PMCPMC4210199.

8. International Stem Cell Initiative C, Akopian V, Andrews PW, Beil S, Benvenisty N, Brehm J, et al. Comparison of defined culture systems for feeder cell free propagation of human embryonic stem cells. In Vitro Cell Dev Biol Anim. 2010;46(3-4):247–58. Epub 2010/02/27. doi: 10.1007/s11626-010-9297-z. PubMed PMID: 20186512; PubMed Central PMCID: PMCPMC2855804.

9. Bai Q, Ramirez JM, Becker F, Pantesco V, Lavabre-Bertrand T, Hovatta O, et al. Temporal analysis of genome alterations induced by single-cell passaging in human embryonic stem cells. Stem Cells Dev. 2015;24(5):653–62. Epub 2014/09/26. doi: 10.1089/scd.2014.0292. PubMed PMID: 25254421; PubMed Central PMCID: PMCPMC4333508.

10. Fu Y, Foden JA, Khayter C, Maeder ML, Reyon D, Joung JK, et al. High-frequency off-target mutagenesis induced by CRISPR-Cas nucleases in human cells. Nat Biotechnol. 2013;31(9):822–6. Epub 2013/06/25. doi: 10.1038/nbt.2623. PubMed PMID: 23792628; PubMed Central PMCID: PMCPMC3773023.

11. Yang L, Grishin D, Wang G, Aach J, Zhang CZ, Chari R, et al. Targeted and genome-wide sequencing reveal single nucleotide variations impacting specificity of Cas9 in human stem cells. Nat Commun. 2014;5:5507. Epub 2014/11/27. doi: 10.1038/ncomms6507. PubMed PMID: 25425480; PubMed Central PMCID: PMCPMC4352754.

12. Laurent LC, Ulitsky I, Slavin I, Tran H, Schork A, Morey R, et al. Dynamic changes in the copy number of pluripotency and cell proliferation genes in human ESCs and iPSCs during reprogramming and time in culture. Cell Stem Cell. 2011;8(1):106–18. Epub 2011/01/08. doi: 10.1016/j.stem.2010.12.003. PubMed PMID: 21211785; PubMed Central PMCID: PMCPMC3043464.

13. Peterson SE, Westra JW, Rehen SK, Young H, Bushman DM, Paczkowski CM, et al. Normal human pluripotent stem cell lines exhibit pervasive mosaic aneuploidy. PLoS One. 2011;6(8):e23018. Epub 2011/08/23. doi: 10.1371/journal.pone.0023018. PubMed PMID: 21857983; PubMed Central PMCID: PMCPMC3156708.

14. Taapken SM, Nisler BS, Newton MA, Sampsell-Barron TL, Leonhard KA, McIntire EM, et al. Karotypic abnormalities in human induced pluripotent stem cells and embryonic stem cells. Nat Biotechnol. 2011;29(4):313–4. Epub 2011/04/12. doi: 10.1038/nbt.1835. PubMed PMID: 21478842.

15. Martin U. Genome stability of programmed stem cell products. Adv Drug Deliv Rev. 2017;120:108–17. Epub 2017/09/18. doi: 10.1016/j.addr.2017.09.004. PubMed PMID: 28917518.

16. Engle SJ, Blaha L, Kleiman RJ. Best Practices for Translational Disease Modeling Using Human iPSC-Derived Neurons. Neuron. 2018;100(4):783–97. Epub 2018/11/23. doi: 10.1016/j.neuron.2018.10.033. PubMed PMID: 30465765.

17. Rohani L, Johnson AA, Naghsh P, Rancourt DE, Ulrich H, Holland H. Concise Review: Molecular Cytogenetics and Quality Control: Clinical Guardians for Pluripotent Stem Cells. Stem Cells Transl Med. 2018;7(12):867–75. Epub 2018/09/16. doi: 10.1002/sctm.18-0087. PubMed PMID: 30218497; PubMed Central PMCID: PMCPMC6265634.

18. Assou S, Bouckenheimer J, De Vos J. Concise Review: Assessing the Genome Integrity of Human Induced Pluripotent Stem Cells: What Quality Control Metrics? Stem Cells. 2018;36(6):814–21. Epub 2018/02/15. doi: 10.1002/stem.2797. PubMed PMID: 29441649.

19. Attwood SW, Edel MJ. iPS-Cell Technology and the Problem of Genetic Instability-Can It Ever Be Safe for Clinical Use? J Clin Med. 2019;8(3). Epub 2019/03/03. doi: 10.3390/jcm8030288. PubMed PMID: 30823421; PubMed Central PMCID: PMCPMC6462964.

20. Baker D, Hirst AJ, Gokhale PJ, Juarez MA, Williams S, Wheeler M, et al. Detecting Genetic Mosaicism in Cultures of Human Pluripotent Stem Cells. Stem Cell Reports. 2016;7(5):998–1012. Epub 2016/11/10. doi: 10.1016/j.stemcr.2016.10.003. PubMed PMID: 27829140; PubMed Central PMCID: PMCPMC5106530.

21. Ben-David U, Mayshar Y, Benvenisty N. Large-scale analysis reveals acquisition of lineage-specific chromosomal aberrations in human adult stem cells. Cell Stem Cell. 2011;9(2):97–102. Epub 2011/08/06. doi: 10.1016/j.stem.2011.06.013. PubMed PMID: 21816361.

22. Israel MA, Yuan SH, Bardy C, Reyna SM, Mu Y, Herrera C, et al. Probing sporadic and familial Alzheimer’s disease using induced pluripotent stem cells. Nature. 2012;482(7384):216–20. Epub 2012/01/27. doi: 10.1038/nature10821. PubMed PMID: 22278060; PubMed Central PMCID: PMCPMC3338985.

23. Chambers SM, Fasano CA, Papapetrou EP, Tomishima M, Sadelain M, Studer L. Highly efficient neural conversion of human ES and iPS cells by dual inhibition of SMAD signaling. Nat Biotechnol. 2009;27(3):275–80. Epub 2009/03/03. doi: 10.1038/nbt.1529. PubMed PMID: 19252484; PubMed Central PMCID: PMCPmc2756723.

24. Wen W, Zhang JP, Xu J, Su RJ, Neises A, Ji GZ, et al. Enhanced Generation of Integration-free iPSCs from Human Adult Peripheral Blood Mononuclear Cells with an Optimal Combination of Episomal Vectors. Stem Cell Reports. 2016;6(6):873–84. Epub 2016/05/11. doi: 10.1016/j.stemcr.2016.04.005. PubMed PMID: 27161365; PubMed Central PMCID: PMCPMC4911493.

25. Feoktistova M, Geserick P, Leverkus M. Crystal Violet Assay for Determining Viability of Cultured Cells. Cold Spring Harb Protoc. 2016;2016(4):pdb prot087379. Epub 2016/04/03. doi: 10.1101/pdb.prot087379. PubMed PMID: 27037069.

26. Zhang SC, Wernig M, Duncan ID, Brustle O, Thomson JA. In vitro differentiation of transplantable neural precursors from human embryonic stem cells. Nat Biotechnol. 2001;19(12):1129–33. Epub 2001/12/04. doi: 10.1038/nbt1201-1129. PubMed PMID: 11731781.

27. Takahashi K, Tanabe K, Ohnuki M, Narita M, Ichisaka T, Tomoda K, et al. Induction of pluripotent stem cells from adult human fibroblasts by defined factors. Cell. 2007;131(5):861–72. doi: 10.1016/j.cell.2007.11.019. PubMed PMID: 18035408.

28. Maherali N, Sridharan R, Xie W, Utikal J, Eminli S, Arnold K, et al. Directly reprogrammed fibroblasts show global epigenetic remodeling and widespread tissue contribution. Cell Stem Cell. 2007;1(1):55–70. Epub 2008/03/29. doi: 10.1016/j.stem.2007.05.014. PubMed PMID: 18371336.

29. Mikkelsen TS, Hanna J, Zhang X, Ku M, Wernig M, Schorderet P, et al. Dissecting direct reprogramming through integrative genomic analysis. Nature. 2008;454(7200):49–55. Epub 2008/05/30. doi: 10.1038/nature07056. PubMed PMID: 18509334; PubMed Central PMCID: PMCPMC2754827.

30. Okita K, Ichisaka T, Yamanaka S. Generation of germline-competent induced pluripotent stem cells. Nature. 2007;448(7151):313–7. Epub 2007/06/08. doi: 10.1038/nature05934. PubMed PMID: 17554338.

31. Wernig M, Meissner A, Foreman R, Brambrink T, Ku M, Hochedlinger K, et al. In vitro reprogramming of fibroblasts into a pluripotent ES-cell-like state. Nature. 2007;448(7151):318–24. Epub 2007/06/08. doi: 10.1038/nature05944. PubMed PMID: 17554336.

32. Allegrucci C, Young LE. Differences between human embryonic stem cell lines. Hum Reprod Update. 2007;13(2):103–20. Epub 2006/08/29. doi: 10.1093/humupd/dml041. PubMed PMID: 16936306.

33. Son MY, Choi H, Han YM, Cho YS. Unveiling the critical role of REX1 in the regulation of human stem cell pluripotency. Stem Cells. 2013;31(11):2374–87. Epub 2013/08/14. doi: 10.1002/stem.1509. PubMed PMID: 23939908.

34. Lowry WE, Richter L, Yachechko R, Pyle AD, Tchieu J, Sridharan R, et al. Generation of human induced pluripotent stem cells from dermal fibroblasts. Proc Natl Acad Sci U S A. 2008;105(8):2883–8. Epub 2008/02/22. doi: 10.1073/pnas.0711983105. PubMed PMID: 18287077; PubMed Central PMCID: PMCPMC2268554.

35. Chin MH, Mason MJ, Xie W, Volinia S, Singer M, Peterson C, et al. Induced pluripotent stem cells and embryonic stem cells are distinguished by gene expression signatures. Cell Stem Cell. 2009;5(1):111–23. Epub 2009/07/03. doi: 10.1016/j.stem.2009.06.008. PubMed PMID: 19570518; PubMed Central PMCID: PMCPMC3448781.

36. Cabrera CM, Cobo F, Nieto A, Cortes JL, Montes RM, Catalina P, et al. Identity tests: determination of cell line cross-contamination. Cytotechnology. 2006;51(2):45–50. Epub 2008/11/13. doi: 10.1007/s10616-006-9013-8. PubMed PMID: 19002894; PubMed Central PMCID: PMCPMC3449683.

37. Chatterjee R. Cell biology. Cases of mistaken identity. Science. 2007;315(5814):928–31. Epub 2007/02/17. doi: 10.1126/science.315.5814.928. PubMed PMID: 17303729.

38. Yaffe MP, Noggle SA, Solomon SL. Raising the standards of stem cell line quality. Nat Cell Biol. 2016;18(3):236–7. Epub 2016/02/26. doi: 10.1038/ncb3313. PubMed PMID: 26911906.

39. Amps K, Andrews PW, Anyfantis G, Armstrong L, Avery S, Baharvand H, et al. Screening ethnically diverse human embryonic stem cells identifies a chromosome 20 minimal amplicon conferring growth advantage. Nature Biotechnology. 2011;29(12):1132–44. doi: 10.1038/nbt.2051.

40. Lund RJ, Narva E, Lahesmaa R. Genetic and epigenetic stability of human pluripotent stem cells. Nat Rev Genet. 2012;13(10):732–44. Epub 2012/09/12. doi: 10.1038/nrg3271. PubMed PMID: 22965355.

41. Hussein SM, Batada NN, Vuoristo S, Ching RW, Autio R, Narva E, et al. Copy number variation and selection during reprogramming to pluripotency. Nature. 2011;471(7336):58–62. Epub 2011/03/04. doi: 10.1038/nature09871. PubMed PMID: 21368824.

42. Narva E, Autio R, Rahkonen N, Kong L, Harrison N, Kitsberg D, et al. High-resolution DNA analysis of human embryonic stem cell lines reveals culture-induced copy number changes and loss of heterozygosity. Nat Biotechnol. 2010;28(4):371–7. Epub 2010/03/31. doi: 10.1038/nbt.1615. PubMed PMID: 20351689.

43. Draper JS, Smith K, Gokhale P, Moore HD, Maltby E, Johnson J, et al. Recurrent gain of chromosomes 17q and 12 in cultured human embryonic stem cells. Nat Biotechnol. 2004;22(1):53–4. Epub 2003/12/09. doi: 10.1038/nbt922. PubMed PMID: 14661028.

44. Inzunza J, Sahlen S, Holmberg K, Stromberg AM, Teerijoki H, Blennow E, et al. Comparative genomic hybridization and karyotyping of human embryonic stem cells reveals the occurrence of an isodicentric X chromosome after long-term cultivation. Mol Hum Reprod. 2004;10(6):461–6. Epub 2004/03/27. doi: 10.1093/molehr/gah051. PubMed PMID: 15044603.

45. Lefort N, Feyeux M, Bas C, Feraud O, Bennaceur-Griscelli A, Tachdjian G, et al. Human embryonic stem cells reveal recurrent genomic instability at 20q11.21. Nat Biotechnol. 2008;26(12):1364–6. Epub 2008/11/26. doi: 10.1038/nbt.1509. PubMed PMID: 19029913.

46. Sareen D, McMillan E, Ebert AD, Shelley BC, Johnson JA, Meisner LF, et al. Chromosome 7 and 19 trisomy in cultured human neural progenitor cells. PLoS One. 2009;4(10):e7630. Epub 2009/11/10. doi: 10.1371/journal.pone.0007630. PubMed PMID: 19898616; PubMed Central PMCID: PMCPMC2765070.

47. Zeng H, Guo M, Martins-Taylor K, Wang X, Zhang Z, Park JW, et al. Specification of region-specific neurons including forebrain glutamatergic neurons from human induced pluripotent stem cells. PLoS One. 2010;5(7):e11853. Epub 2010/08/06. doi: 10.1371/journal.pone.0011853. PubMed PMID: 20686615; PubMed Central PMCID: PMCPMC2912324.

48. Lamm N, Ben-David U, Golan-Lev T, Storchova Z, Benvenisty N, Kerem B. Genomic Instability in Human Pluripotent Stem Cells Arises from Replicative Stress and Chromosome Condensation Defects. Cell Stem Cell. 2016;18(2):253–61. Epub 2015/12/17. doi: 10.1016/j.stem.2015.11.003. PubMed PMID: 26669899.

49. Ruiz S, Lopez-Contreras AJ, Gabut M, Marion RM, Gutierrez-Martinez P, Bua S, et al. Limiting replication stress during somatic cell reprogramming reduces genomic instability in induced pluripotent stem cells. Nat Commun. 2015;6:8036. Epub 2015/08/22. doi: 10.1038/ncomms9036. PubMed PMID: 26292731; PubMed Central PMCID: PMCPMC4560784.

50. O’Malley J, Skylaki S, Iwabuchi KA, Chantzoura E, Ruetz T, Johnsson A, et al. High-resolution analysis with novel cell-surface markers identifies routes to iPS cells. Nature. 2013;499(7456):88–91. Epub 2013/06/04. doi: 10.1038/nature12243. PubMed PMID: 23728301; PubMed Central PMCID: PMCPMC3743022.

51. Papapetrou EP, Tomishima MJ, Chambers SM, Mica Y, Reed E, Menon J, et al. Stoichiometric and temporal requirements of Oct4, Sox2, Klf4, and c-Myc expression for efficient human iPSC induction and differentiation. Proc Natl Acad Sci U S A. 2009;106(31):12759–64. Epub 2009/06/25. doi: 10.1073/pnas.0904825106. PubMed PMID: 19549847; PubMed Central PMCID: PMCPMC2722286.

52. Yamaguchi S, Hirano K, Nagata S, Tada T. Sox2 expression effects on direct reprogramming efficiency as determined by alternative somatic cell fate. Stem Cell Res. 2011;6(2):177–86. Epub 2010/12/07. doi: 10.1016/j.scr.2010.09.004. PubMed PMID: 21130722.

53. Matoba R, Niwa H, Masui S, Ohtsuka S, Carter MG, Sharov AA, et al. Dissecting Oct3/4-regulated gene networks in embryonic stem cells by expression profiling. PLoS One. 2006;1:e26. Epub 2006/12/22. doi: 10.1371/journal.pone.0000026. PubMed PMID: 17183653; PubMed Central PMCID: PMCPMC1762406.

54. Daniszewski M, Nguyen Q, Chy HS, Singh V, Crombie DE, Kulkarni T, et al. Single-Cell Profiling Identifies Key Pathways Expressed by iPSCs Cultured in Different Commercial Media. iScience. 2018;7:30–9. Epub 2018/09/30. doi: 10.1016/j.isci.2018.08.016. PubMed PMID: 30267684; PubMed Central PMCID: PMCPMC6135898.

55. Nguyen QH, Lukowski SW, Chiu HS, Senabouth A, Bruxner TJC, Christ AN, et al. Single-cell RNA-seq of human induced pluripotent stem cells reveals cellular heterogeneity and cell state transitions between subpopulations. Genome Res. 2018;28(7):1053–66. Epub 2018/05/13. doi: 10.1101/gr.223925.117. PubMed PMID: 29752298; PubMed Central PMCID: PMCPMC6028138.

56. Koehler KR, Tropel P, Theile JW, Kondo T, Cummins TR, Viville S, et al. Extended passaging increases the efficiency of neural differentiation from induced pluripotent stem cells. BMC Neurosci. 2011;12:82. Epub 2011/08/13. doi: 10.1186/1471-2202-12-82. PubMed PMID: 21831300; PubMed Central PMCID: PMCPMC3167757.

57. Yu J, Vodyanik MA, Smuga-Otto K, Antosiewicz-Bourget J, Frane JL, Tian S, et al. Induced pluripotent stem cell lines derived from human somatic cells. Science. 2007;318(5858):1917–20. doi: 10.1126/science.1151526. PubMed PMID: 18029452.

58. Peterson SE, Tran HT, Garitaonandia I, Han S, Nickey KS, Leonardo T, et al. Teratoma generation in the testis capsule. J Vis Exp. 2011;(57):e3177. Epub 2011/12/14. doi: 10.3791/3177. PubMed PMID: 22158256; PubMed Central PMCID: PMCPMC3308584.

59. Muller FJ, Brandl B, Loring JF. Assessment of human pluripotent stem cells with PluriTest. StemBook. Cambridge (MA)2008.

60. Ungrin MD, Joshi C, Nica A, Bauwens C, Zandstra PW. Reproducible, ultra high-throughput formation of multicellular organization from single cell suspension-derived human embryonic stem cell aggregates. PLoS One. 2008;3(2):e1565. Epub 2008/02/14. doi: 10.1371/journal.pone.0001565. PubMed PMID: 18270562; PubMed Central PMCID: PMCPMC2215775.

61. Itskovitz-Eldor J, Schuldiner M, Karsenti D, Eden A, Yanuka O, Amit M, et al. Differentiation of human embryonic stem cells into embryoid bodies compromising the three embryonic germ layers. Mol Med. 2000;6(2):88–95. Epub 2000/06/20. PubMed PMID: 10859025; PubMed Central PMCID: PMCPMC1949933.

62. Mandai M, Watanabe A, Kurimoto Y, Hirami Y, Morinaga C, Daimon T, et al. Autologous Induced Stem-Cell-Derived Retinal Cells for Macular Degeneration. N Engl J Med. 2017;376(11):1038–46. Epub 2017/03/16. doi: 10.1056/NEJMoa1608368. PubMed PMID: 28296613.

63. Scotto L, Narayan G, Nandula SV, Arias-Pulido H, Subramaniyam S, Schneider A, et al. Identification of copy number gain and overexpressed genes on chromosome arm 20q by an integrative genomic approach in cervical cancer: potential role in progression. Genes Chromosomes Cancer. 2008;47(9):755–65. Epub 2008/05/29. doi: 10.1002/gcc.20577. PubMed PMID: 18506748.

64. Liang G, Zhang Y. Genetic and epigenetic variations in iPSCs: potential causes and implications for application. Cell Stem Cell. 2013;13(2):149–59. Epub 2013/08/06. doi: 10.1016/j.stem.2013.07.001. PubMed PMID: 23910082; PubMed Central PMCID: PMCPMC3760008.

65. Yoshihara M, Hayashizaki Y, Murakawa Y. Genomic Instability of iPSCs: Challenges Towards Their Clinical Applications. Stem Cell Rev Rep. 2017;13(1):7–16. Epub 2016/09/07. doi: 10.1007/s12015-016-9680-6. PubMed PMID: 27592701; PubMed Central PMCID: PMCPMC5346115.

66. Xu RH, Sampsell-Barron TL, Gu F, Root S, Peck RM, Pan G, et al. NANOG is a direct target of TGFbeta/activin-mediated SMAD signaling in human ESCs. Cell Stem Cell. 2008;3(2):196–206. Epub 2008/08/07. doi: 10.1016/j.stem.2008.07.001. PubMed PMID: 18682241; PubMed Central PMCID: PMCPMC2758041.

67. D’Amour KA, Agulnick AD, Eliazer S, Kelly OG, Kroon E, Baetge EE. Efficient differentiation of human embryonic stem cells to definitive endoderm. Nat Biotechnol. 2005;23(12):1534–41. Epub 2005/11/01. doi: 10.1038/nbt1163. PubMed PMID: 16258519.

68. Xu RH, Chen X, Li DS, Li R, Addicks GC, Glennon C, et al. BMP4 initiates human embryonic stem cell differentiation to trophoblast. Nat Biotechnol. 2002;20(12):1261–4. Epub 2002/11/12. doi: 10.1038/nbt761. PubMed PMID: 12426580.

